# The histidine kinase VraS orchestrates the cell-wall stress response in *Staphylococcus aureus* via its direct interaction with glycopeptides and β-lactams

**DOI:** 10.1101/2025.01.23.634444

**Authors:** Melisa B. Antinori, David Sychantha, Kalinka Koteva, Irina P. Suarez, Melani D. Peralta, Gerard D. Wright, Leticia I. Llarrull

## Abstract

Multidrug-resistant *Staphylococcus aureus* is a major global health threat, with the VraTSR three-component system playing a key role in sensing and conferring resistance to cell-wall active antibiotics, particularly vancomycin. VraTSR comprises the membrane histidine kinase VraS, the cytoplasmic response regulator VraR, and the uncharacterized membrane protein VraT, which regulate the cell wall stress stimulon. However, the molecular signals sensed by VraTSR remain unknown. To elucidate the activation mechanism of this regulatory system, we investigated interactions with β-lactams and glycopeptides. Using a transcriptional reporter strain, we confirmed VraTSR activation by β-lactams, glycopeptides, a vancomycin-derived photoprobe (VPP), and the previously unreported activators A47934 and moenomycin A. Photo-crosslinking assays with VPP and full-length VraS expressed in membranes revealed a direct interaction with vancomycin, which was further confirmed in purified VraS reconstituted in liposomes. VPP binding was concentration-dependent, saturable, and displaced by vancomycin. Saturation transfer difference (STD) Nuclear Magnetic Resonance (NMR) experiments confirmed vancomycin binding to VraS and demonstrated ampicillin interaction, highlighting the involvement of aryl protons from both antibiotics. These findings establish VraS as a receptor for vancomycin and ampicillin. In contrast, assays with membrane vesicles expressing only VraT or co-expressing VraS/VraT did not show covalent adduct formation between VraT and VPP. While VraT’s exact role remains unclear, its participation in antibiotic sensing or signal transduction cannot yet be excluded. These results demonstrate that vancomycin and ampicillin directly activate VraS, providing critical insights into the activation of the cell wall stress stimulon and the mechanisms underlying antibiotic resistance. Disrupting VraTSR signaling is a promising strategy to combat multidrug resistance in *S. aureus,* and we provide invaluable *in vitro* platforms for identifying potential VraS inhibitors.

**Author Summary:** Multidrug-resistant *Staphylococcus aureus* poses a major global health threat due to its resistance to cell-wall active antibiotics. Our study focuses on the VraTSR three-component system, a key regulator of the cell wall stress response in *S. aureus*, whose activation signals have remained unknown.

We demonstrate that VraS, the membrane histidine kinase of the system, acts as a direct receptor for vancomycin and ampicillin—two structurally distinct antibiotics. These findings uncover the activation mechanism of VraTSR and position VraS as a central player in antibiotic sensing and resistance.

By identifying VraS as a direct antibiotic receptor, we provide a promising target for developing inhibitors to disrupt VraTSR signaling and restore antibiotic efficacy. Additionally, the *in vitro* platforms we established enable the identification and testing of potential VraS inhibitors.

This study highlights the importance of understanding bacterial stress-response pathways to combat antibiotic resistance, offering critical insights for developing new therapeutic strategies against multidrug-resistant *S. aureus*, a growing global health challenge.

## Introduction

*Staphylococcus aureus* is the leading cause of hospital-and community-associated infections [1–3] often linked to lower respiratory tract and cardiovascular infections. In addition, it is the second most common cause of healthcare-associated pneumonia and bloodstream infections [1, 2, 4, 5]. Methicillin-resistant *S. aureus* (MRSA) is a high-priority global concern in hospitals and the community [6]. MRSA has become the first cause of hospital-associated infections in Latin America, and there has been an increasing number of reports of community-acquired MRSA infections [7, 8]. The acquisition of resistance to β-lactam antibiotics, generally concurrent with the acquisition of resistance to other antibacterial agents, represents a considerable challenge for preventing and treating *S. aureus-*associated infections [9–11].

The VraTSR cell-wall-stress regulatory system of *S. aureus* is implicated in coordinating important steps of cell wall biosynthesis and maintaining the integrity of the peptidoglycan [12, 13]. Activation of VraTSR triggers the transcription of a group of genes called the Cell Wall Stress Stimulon (CWSS), which upregulates cell wall synthesis, causing it to thicken [13–16]. Thickening of the cell wall gives rise to the vancomycin-intermediate resistance phenotype in *S. aureus* (VISA strains). The VraTSR system is induced by antibiotics that interfere with different stages of cell wall synthesis, including β-lactams, glycopeptides, daptomycin, bacitracin, tunicamycin, D-cycloserine, fosfomycin, flavomycin, teicoplanin, and [2-(2-chlorophenyl)-3-[1-(2,3-dimethylbenzyl) piperidin-4-yl]-5-fluoro-1H-indole] (CDFI), an inhibitor of a protein with predicted peptidoglycan lipid II flippase function [16, 17]. Mutations within this system have been shown to lead to susceptibility to some of these inducers, namely glycopeptides, β-lactams, bacitracin, and D-cycloserine [18–20]. Moreover, the inactivation of VraTSR abolishes β-lactam resistance in a variety of genetic backgrounds that are phenotypically β-lactam-resistant, regardless of the presence of *mecI* and SCCmec types II or IV in MRSA [21–24]. Despite being natively encoded in the *S. aureus* chromosome, VraTSR can enhance the expression of horizontally acquired enterococcal *vanA* gene cluster, increasing resistance to glycopeptides [25].

The VraTSR system shares homology with other well-studied systems, such as the LiaRS system from *Bacillus subtilis*, *Streptococcus pneumoniae*, and *Streptococcus mutans* [26, 27] or the CesSR system from *Lactococcus* species [28]. The genes that belong to this family are co-transcribed with a third gene with an accessory regulatory function, which is essential for the activation of the system and survival of the bacterium [26, 29]. Hence, instead of two-component systems (TCS), they constitute three-component systems. The VraTSR operon comprises the *vraU*, *vraT*, *vraS,* and *vraR* genes (Fig 1), where the last three encode the VraT, VraS, and VraR proteins, respectively, which are common to other regulatory systems [30]. VraS is the histidine kinase, and VraR is the system’s response regulator. A mutant lacking the *vraU* gene is phenotypically identical to the wild-type strain, indicating that *vraU* is dispensable for the induction of the system or the activation of the CWSS [30]. In contrast, disruption of any of the other three genes prevents activation of the VraTSR system, rendering the bacterium susceptible to β-lactam antibiotics and vancomycin [19, 30–32]. VraT is a membrane protein with a LiaF-like extracellular domain of unknown function required for optimal expression of the resistant phenotype [33].

**Fig 1.**
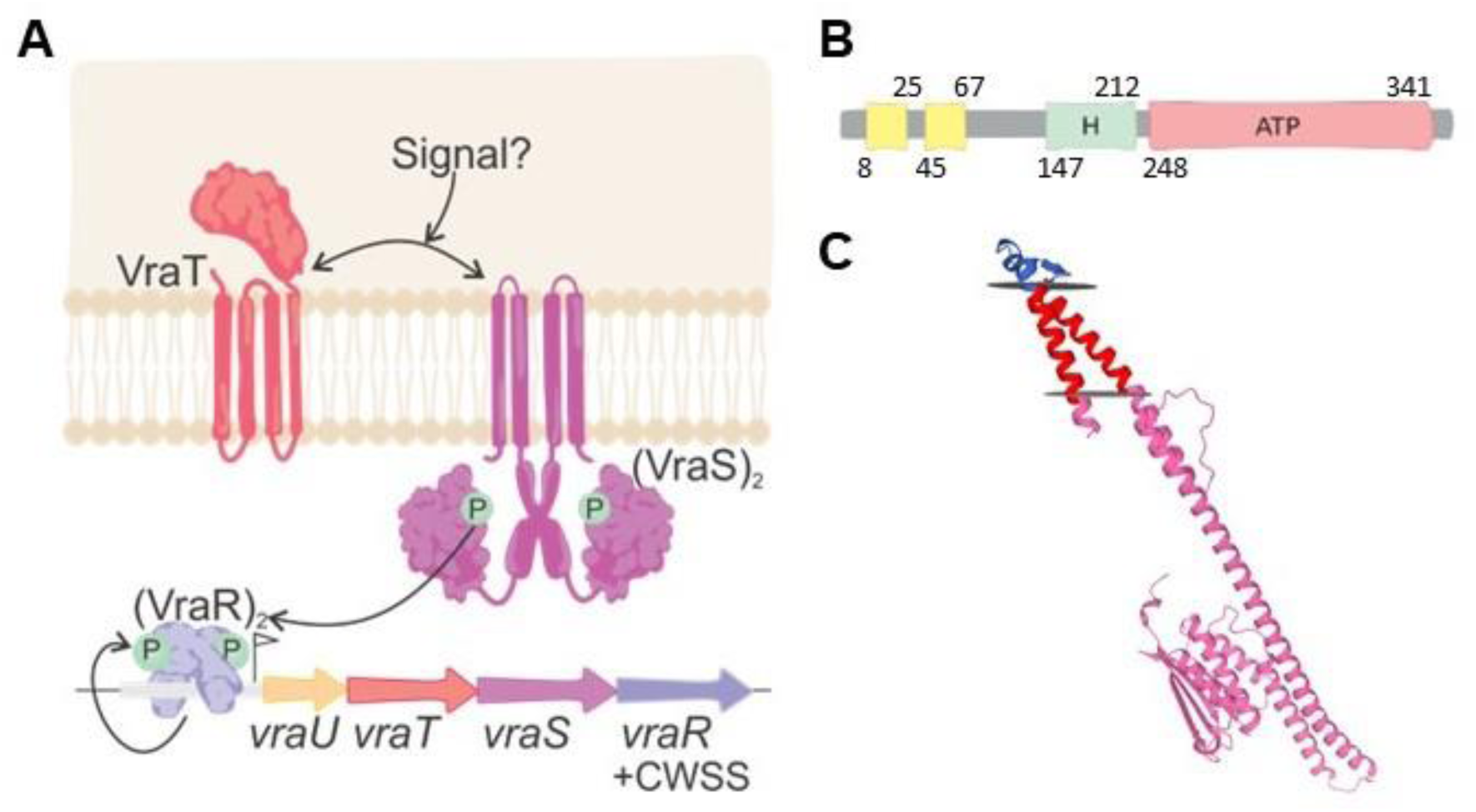
Schematic for the activation of the VraTSR system and predicted topology of VraS. **A.** Recognition of the presence of glycopeptide or β-lactam antibiotics in the extracellular space is proposed to occur either through the extracellular loop of VraS and/or the extracellular domain of VraT. Activation of VraS leads to autophosphorylation, followed by phosphotransfer to the response regulator VraR. Phosphorylated VraR activates transcription of the genes of the VraTSR operon and triggers the transcription of a group of genes called the Cell Wall Stress Stimulon (CWSS) and to the vancomycin-intermediate phenotype in *S. aureus*. **B.** Predicted topology of VraS (DeepTMHMM). Yellow: transmembrane α-helices. Residues 68 to 347: the C-terminal cytoplasmic core of VraS, containing the dimerization domain (with the conserved His^156^ residue that is phosphorylated) and the ATP-binding domain (residues 248 to 341). **C.** Transmembrane protein topology prediction for VraS in combination with the predicted protein structure for the VraS monomer. The predicted extracellular loop (residues 26-44; blue), the two transmembrane α-helices (residues 8-25 and 48-68; red), and the C-terminal cytoplasmic domain (with the catalytic His residue, the dimerization, and the ATP-binding domains). The crystal structure of residues 212 to 341 comprising the ATP-binding domain has been reported (PDB 4GT8Z).

The mechanism of VraTSR activation by cell-wall active antibiotics is unknown. Considering that both VraS and VraT possess extracellular segments, we hypothesize that the VraTSR system could be triggered by a molecular signal that interacts with these regions [33]. As diverse antibiotics induce the VraTSR system, one possibility is that VraTSR is activated by cell wall degradation products or intermediates of peptidoglycan synthesis. Peptidoglycan fragments or intermediates have been reported to regulate enzymes and transcription repressors in several systems [34–37]. Alternatively, antibiotics may interact directly with the VraTSR membrane proteins. VraS shares homology with the VanS histidine kinase of the VanSR system of *Streptomyces ceolicolor*, which has been shown to interact with vancomycin directly with its extracellular region [38–41].

To identify the molecular signal sensed by the VraTSR system, we investigated its activation by a panel of antibiotics and peptidoglycan fragments. We showed activation of the VraTSR system could not be achieved with PG fragments but was robustly induced by cell wall-targeting antibiotics, including a new class of cell-wall active antibiotics: phosphoglycolipids. In addition, using a vancomycin-derived photoprobe and saturation transfer difference (STD) NMR experiments, we demonstrated the direct interaction of the glycopeptide antibiotic vancomycin with VraS, an interaction that did not depend on the presence of VraT. The STD experiments also provided evidence of ampicillin binding to purified full-length VraS. Furthermore, the STD assays showed that aromatic groups of vancomycin and ampicillin were involved in the interaction with VraS.

## Results

### Activation of the VraTSR system by cell-wall active antibiotics and peptidoglycan fragments

The effect of cell-wall active antibiotics was evaluated using an *S. aureus* RN4220 reporter strain. We worked with *S. aureus* RN4220 cells −which harbor a chromosomal copy of the *VraTSR* operon− transformed with the pCN52::*opvra* plasmid. In this reporter strain, the *gfp* gene is under the control of the operator region of the VraTSR system. Incubation of the reporter strain with the β-lactam antibiotics ampicillin and oxacillin and with the glycopeptide antibiotic vancomycin resulted in an increase in the intensity of fluorescence due to GFP expression, as expected since it is widely reported that these antibiotics induce the expression of genes of the VraTSR system (Fig 2.A). Vancomycin induced the VraTSR system at concentrations equal to and greater than 2.5 μg/ml (approximately four times the MIC), remaining constant at the highest concentrations used. Ampicillin concentrations lower than 0.5 μg/ml (concentration corresponding to the MIC) did not induce the system. In comparison, ampicillin concentrations equal to and greater than 0.5 μg/ml (corresponding to the MIC and up to twenty times higher) activated the system, reaching similar fluorescence levels. Activation by oxacillin showed a similar behavior: activation was observed starting from a concentration of 0.25 μg/ml (equivalent to the MIC), and using higher concentrations of antibiotic (from two to twenty times the MIC) resulted in higher fluorescence intensity. Our results align with previous reports that activation of the VraTSR system in *S. aureus* reporter strains is seen at vancomycin and oxacillin concentrations greater than the MIC [42, 43]. Based on these results, it was decided to establish the 5 μg/ml (approximately eight times the MIC) as a fixed concentration for future activation assays. This amount of antibiotic ensured the induction of the VraTSR system by all three antibiotics, resulting in the highest relative fluorescence intensity recorded.

**Fig 2.**
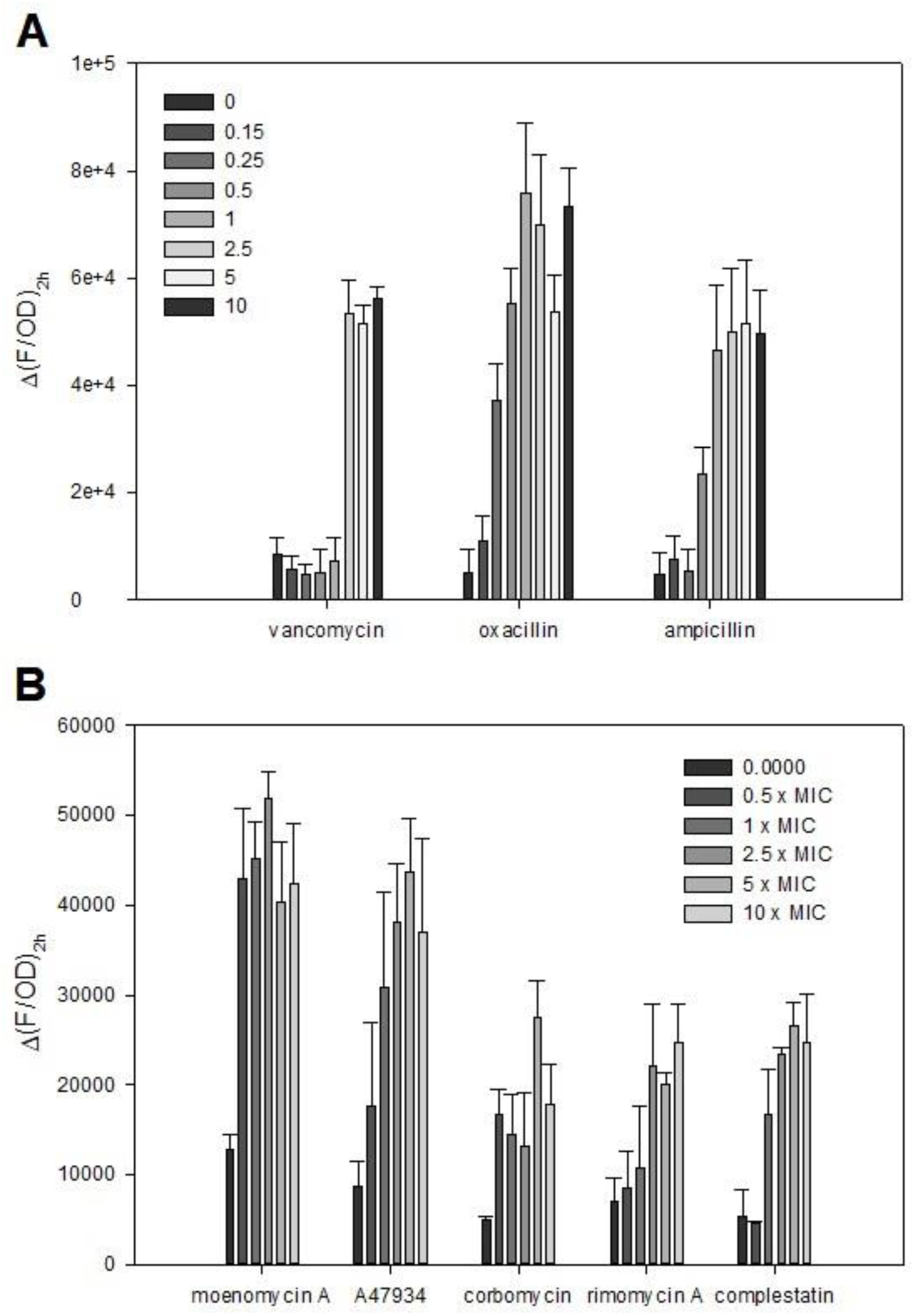
Activation of the VraTSR system by cell-wall active antibiotics. **A.** Activation by β-lactam and glycopeptide antibiotics. Change in relative fluorescence after incubation for 2 h (Δ(F/OD)_2h_) with different concentrations of vancomycin (Van), oxacillin (Oxa), and ampicillin (Amp); the Δ(F/OD)_2h_ is plotted. **B.** Activation by moenomycin A, A47934, corbomycin, rimomycin-A, and complestatin. Change in relative fluorescence after incubation for 2 hours (Δ(F/OD)_2h_) with different concentrations of the indicated antibiotic. In A and B, the reporter strain *S. aureus* RN4220/pCN52::*opvra* and the control strain *S. aureus* RN4220/pCN52 were grown until OD_600nm_ = 0.3, at which point the cultures were aliquoted in 96-well plates. The different antibiotics were added at the indicated concentrations and incubated for 2 h at 37 °C in an Agilent BioTek Synergy Neo2 Hybrid Multi-Mode Microplate Reader. In *S. aureus* RN4220/pCN52::*opvra* cultures, the intensity of GFP fluorescence emission was recorded at λ_em_ = 528 nm, upon excitation at 485 nm and OD_600nm_ was measured simultaneously; at each time point, F/OD_600nm_ was calculated. F/OD_600nm_ was also calculated for the control strain *S. aureus* RN4220/pCN52 (autofluorescence) and was subtracted from the relative fluorescence of the reporter strain *S. aureus* RN4220/pCN52::*opvra*, to give (F/OD)_t_ = (F/ODpCN52::*opvra*)_t_ – (F/ODpCN52)_t_. The graphs show Δ(F/OD)_2h_ = (F/ODpCN52::*opvra* – F/ODpCN52)_t=2h_ – (F/ODpCN52::*opvra* – F/ODpCN52)_t=0h_. The results represent the means of two experiments, and the error bars correspond to the standard deviations (SD).

Vancomycin binds to the D-Ala-D-Ala terminus of the peptidoglycan precursor lipid II [44]. Formation of the stable lipid II-vancomycin complex blocks the peptidoglycan synthesis cycle by reducing the penicillin-binding proteins (PBPs) substrate availability that catalyze transglycosylation and transpeptidation. Hence, we decided to evaluate whether cell-wall active antibiotics with a different mode of action also induce the VraTSR system. We worked with the glycopeptides complestatin, corbomycin, rimomycin-A, A47934, and the phosphoglycolipid monoemycin A (Fig 3). The Type V glycopeptides corbomycin, complestatin, and rimomycin A indirectly inhibit autolysins by binding peptidoglycan [45–47]; they do not inhibit PG biosynthesis but block cell division. A47934, produced by *Streptomyces toyocaensis*, possesses a nonglycosylated heptapeptide core that is sulfated on the phenolic hydroxyl of the N-terminal 4-hydroxy-L-phenylglycine residue and is, in addition, chlorinated at three distinct sites [48, 49]. A47934 retains the D-Ala-D-Ala binding mode of action of canonical glycopeptide antibiotics [49]. The phosphoglycolipid moenomycin A inhibits bacterial cell wall synthesis by binding to the transglycosylases responsible for peptidoglycan carbohydrate chain formation [50–52].

**Fig 3.**
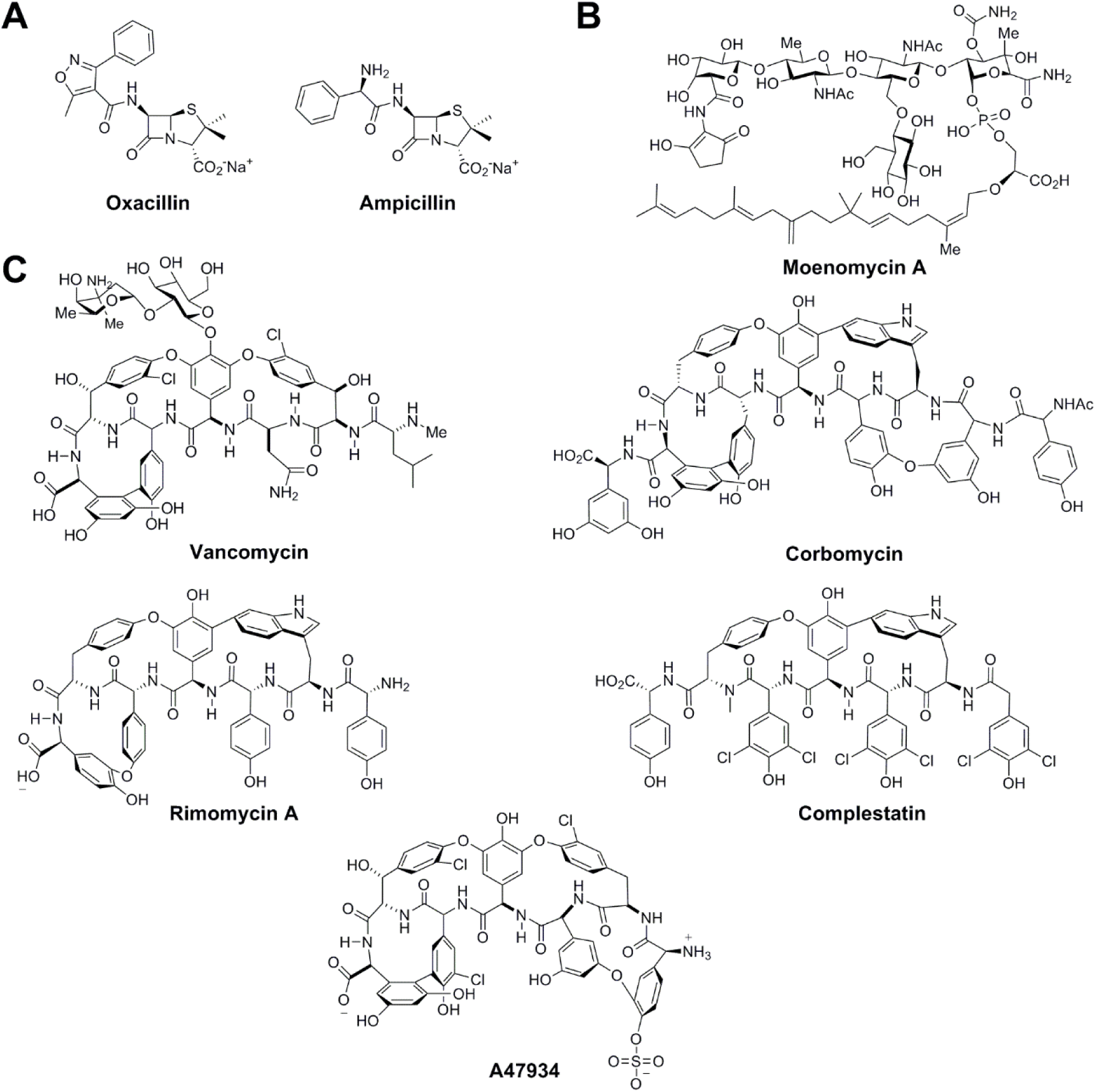
Structures of the antibiotics that activate the VraTSR system and the photoprobes used in this study. **A.** Structure of the β-lactams oxacillin and ampicillin. **B.** Structure of the phosphoglycolipid moenomycin A. **C.** Structure of the glycopeptides vancomycin, corbomycin, rimomycin A, complestatin, and A47934.

The glycopeptides complestatin, corbomycin, rimomycin-A, A47934, and the phosphoglycolipid monoemycin A induced the VraTSR system, although they differed in the activation efficiency (Fig 2.B). Moenomycin A was the strongest inducer and was active at a sub-MIC concentration. However, higher amounts of moenomycin A did not increase the relative fluorescence. Incubation with A47934, for its part, generated a concentration-dependent activation of the system (starting from concentrations 0.5 times the MIC) and reached values similar to those obtained with the reference antibiotics. The Type V glycopeptides rimomycin A and complestatin activated the VraTSR system to a lesser extent, with the maximum fluorescence intensity detected approximately half that recorded for vancomycin. Finally, corbomycin was the weakest inducer, requiring concentrations of 5 times the MIC. The VraTSR system was not induced by antibiotics that do not target the cell wall (S2 Fig). In contrast to published data [53], the lipopeptide daptomycin did not activate the VraTSR system in our assays with the RN4220 reporter strain.

Given that the VraTSR system is activated by cell-wall active antibiotics that differ in structure and mechanism of action, the system may respond to molecules that accumulate upon blockage of cell-wall synthesis, as postulated by Boyle-Vavra and coworkers [30]. To test this hypothesis, we evaluated whether the peptidoglycan fragments generated by the action of mutanolysin and lysostaphin could induce the VraTSR system. At the highest concentration tested, the peptidoglycan fragments generated by these two enzymes alone or in combination did not activate the system (Fig 4). In addition, these enzymatically produced peptidoglycan fragments did not potentiate (or, for that matter, reduce) the effect of ampicillin, oxacillin, and vancomycin.

**Fig 4.**
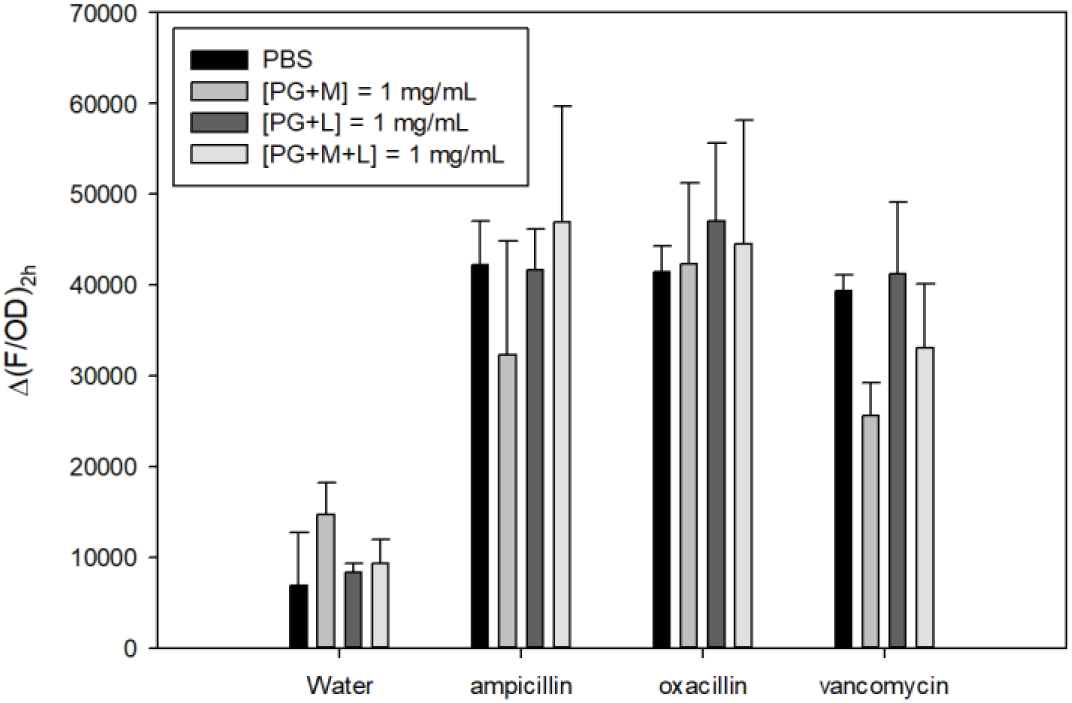
Effect of peptidoglycan fragments and their combination with β-lactam and glycopeptide antibiotics on activating the VraTSR system. Addition of 1 mg/ml of peptidoglycan fragments in the absence or presence of 5 µg/ml of the inducing antibiotic (ampicillin (Amp), oxacillin (Oxa) and vancomycin (Van)). PG+M, fragments generated from the action of mutanolysin on peptidoglycan; PG+L, fragments generated from the action of lysostaphin on peptidoglycan; PG+M+L, fragments generated from the action of both enzymes. The cell cultures were grown until OD_600nm_=0.3, at which point they were aliquoted in 96-well plates, and the different antibiotics, peptidoglycan fragments, or a combination of both were added; cells were incubated for 2 h at 37 °C in an Agilent BioTek Synergy Neo2 Hybrid Multi-Mode Microplate Reader. The graphs show Δ(F/OD)_2h_ = (F/ODpCN52::*opvra* – F/ODpCN52)_t=2h_ – (F/ODpCN52::*opvra* – F/ODpCN52)_t=0h_. The results represent the means of two experiments, and the error bars correspond to the standard deviations (SD).

### A vancomycin-derived photo-affinity probe identifies a direct interaction with VraS

Since vancomycin induces the VraTSR system, generating a vancomycin-intermediate phenotype in *S. aureus*, an affinity photoprobe derived from the glycopeptide antibiotic vancomycin (VPP; Fig 5.A) [40] was used to assess whether it binds to VraS and/or VraT. The VPP includes a biotin label on its C-terminal for visualization and purification of complexes using biotin-specific antibodies or streptavidin/Strep-Tactin and a benzophenone photoreactive group on vancomycin for covalent labeling of interacting proteins. On irradiation with light of 365 nm, a diradical is generated by photolysis at the benzophenone moiety (Fig 5.B) that covalently links to nearby aliphatic amino acids while maintaining antibiotic activity, as described previously [40]. This was confirmed using the VraTSR reporter strain, which showed that the modifications introduced to vancomycin’s structure in generating VPP did not affect its ability to activate VraTSR (S3 Fig).

**Fig 5.**
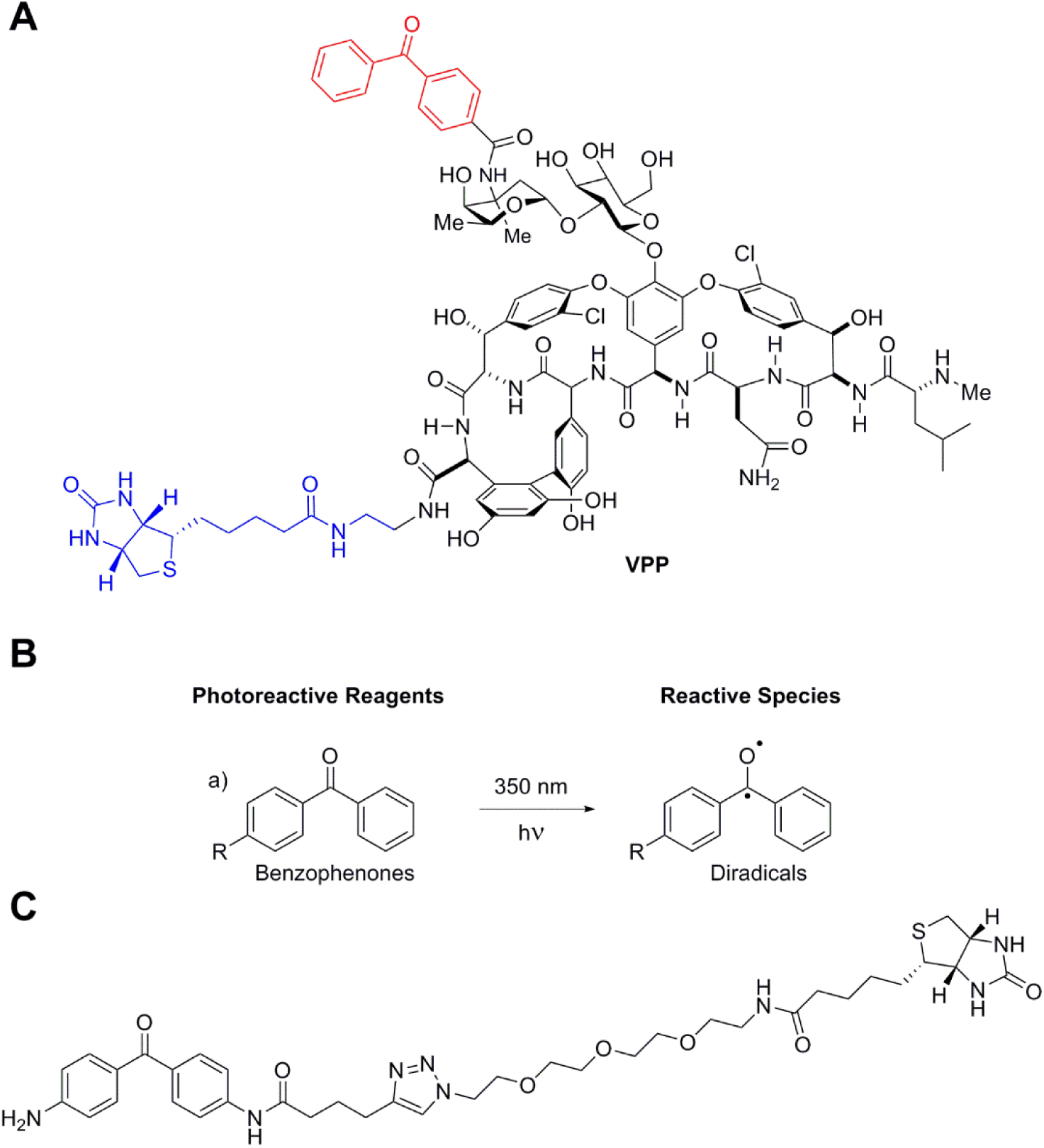
Structures of the photoprobes used in this study. **A.** Structure of the vancomycin-derived photoprobe (VPP); vancomycin backbone (black), biotin label (blue), and benzophenone affinity photoprobe (red). **B.** Benzophenone and active biradical generated from UV irradiation. **C.** Dummy photoprobe, used in this work as a vancomycin specificity control.

To establish whether the activation of the VraTSR system by glycopeptide antibiotics is due to the direct interaction of the antibiotic with the membrane proteins VraS and/or VraT, we overexpressed both proteins in *E. coli* membranes and evaluated the interaction with VPP. Membrane preparations were incubated with VPP, and after irradiation to activate the benzophenone moiety, proteins covalently bound to VPP were detected by Western blot.

In the assay with membranes containing recombinant VraS (VraS-H6x; theoretical mass of 40,867.37 Da), a 40-kDa band was detected in blots developed with biotin-specific and His-tag-specific antibodies, consistent with VPP-bound VraS-H6x (Fig 6.A, red asterisk). Biotinylated VraS-H6x was absent when the same membranes were incubated with excess vancomycin before incubation with VPP and irradiation. Similarly, biotinylated VraS-H6x was not detected in the membranes that were incubated with a control photoprobe (named Dummy; Fig 5.C), consisting of a benzophenone group bound to biotin through a PEG linker. Two additional prominent biotinylated proteins were detected in VraS membranes incubated with VPP. These proteins were also observed, though less distinctly, in membranes from *E. coli* transformed with an empty vector and treated with VPP. The intensity of these bands was reduced upon preincubation with vancomycin. One protein had an approximate mass of 49 kDa (Fig 6, green asterisk), while the other was greater than 62 kDa (Fig 6, blue asterisk). These bands may correspond to native *E. coli* membrane proteins that bind vancomycin, but their identities remain to be determined. The biotinylated band with a mass > 62 kDa (Fig 6, black asterisk) may partially represent residual, non-denatured VraS dimers. In fact, despite denaturing conditions, a > 62 kDa band was detected in Western blots using His-tag-specific antibodies. We confirmed that purified VraS-H6x forms dimers (S4 Fig), consistent with observations for other histidine kinases in TCSs.

**Fig 6.**
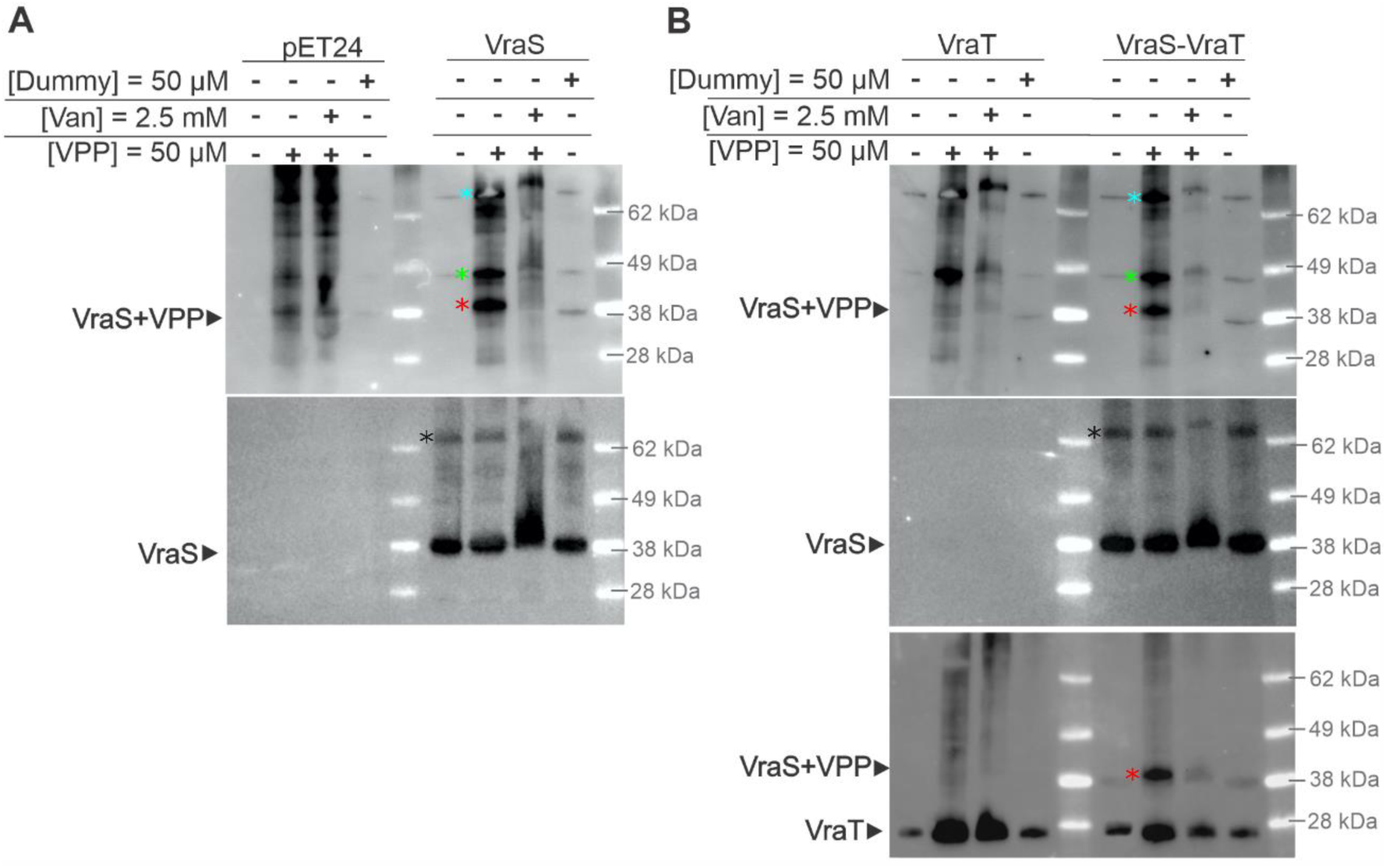
Evaluation of the interaction of the vancomycin-derived photoprobe with VraS and VraT in membrane vesicles. Membrane vesicles in 10 mM HEPES, pH 7, at a total protein concentration of 1 mg/ml, were incubated for 1 h with DMSO, 50 μM Dummy, or 50 μM VPP. In addition, a fraction of membrane vesicles was pre-incubated for 1 h with 2.5 mM vancomycin and then incubated with 50 μM VPP. After incubation with the photoprobes, the preparations were irradiated with UV-light (365 nm) for 2 h at 4 °C. Assays with membrane vesicle preparations from *E. coli* BL21 Star^TM^ (DE3) cells transformed with: **A.** pET24a(+) (control) and pET24a(+)::*vraS-H6x*; **B.** pET24a(+)::*vraT-TST* and pET24a(+)::*vraS-H6x*::*vraT-TST*. In A and B, protein samples (5 µg of total protein in each lane) were resolved by NuPAGE (4-12% gel) and analyzed by Western blot. **In A and B, Top panel:** biotinylated proteins (including VPP-labelled proteins) were detected with a primary antibody specific for biotin (goat α-biotin antibody 1/5,000) and the rabbit α-goat IgG-HRP secondary antibody (1/50,000). *Red asterisk: VraS-VPP adduct. *Green asterisk: putative *E. coli* vancomycin-binding protein crosslinked to VPP. *Blue asterisk: putative *E. coli* vancomycin-binding protein, crosslinked to VPP or VraS dimer bound crosslinked to VPP. **In A, bottom panel and B, middle panel:** VraS-H6x was detected with an antibody specific for the C-terminal His-Tag (Anti-His-Tag-HRP 1/50,000). *Black asterisk: VraS dimer, remaining after denaturation. **In B, bottom panel:** the Twin-Strep-tag in VraT-TST was detected with Streptactin-HRP (1/5,000). The latter reagent also interacts with biotin in VPP; hence, it also detected VPP-labelled VraS. *Red asterisk: VraS-VPP adduct. Sequential chemiluminescent detection was performed on the same membrane by inactivation of HRP by treatment with H_2_O_2_ (in the following order: goat α-biotin antibody; Anti-His-Tag-HRP; Streptactin-HRP). The expected molecular mass for VraS-H6x is 40 kDa, and for VraT-TST is 30 kDa. Assays with VPP and membranes expressing recombinant VraT (VraT-TST) showed no evidence of direct interaction between VPP and VraT (Fig 6.B). A band at approximately 30 kDa, consistent with the expected mass of VraT-TST, was detected in all four lanes. However, no VPP-bound band at ∼30 kDa was observed in Western blots probed with a biotin-specific antibody.

When VraS and VraT were co-expressed, a biotinylated band at approximately 40 kDa (Fig 6.B, red asterisk) was detected by both the biotin-specific antibody and Strep-Tactin-HRP. This band was indistinguishable in mass from the one observed in VraS-expressing membrane vesicles but was absent in samples preincubated with vancomycin before VPP incubation or in those incubated with Dummy.

In summary, these assays with the vancomycin-derived photoprobe and the VraS and VraT proteins in *E. coli* membranes provided the first evidence of direct interaction of vancomycin with VraS, both in the absence and presence of VraT. They also indicated no direct interaction of vancomycin with VraT, at least not one that forms a covalent adduct with the photoprobe used in this work.

### Characterization of the interaction of VraS with the vancomycin-derived photoprobe

Given the evidence of the interaction of vancomycin with VraS expressed in membrane vesicles, we further characterized vancomycin binding to VraS *in vitro*. Full-length VraS was overexpressed in *E. coli* membranes to produce active protein for biochemical assays. Recombinant VraS was purified, reconstituted in DDM micelles, and incorporated into liposomes. Both purified VraS in DDM micelles and proteoliposomes were properly folded and active as judged by circular dichroism and autophosphorylation assays, respectively (Fig 7). Furthermore, purified VraS in DDM micelles and proteoliposomes formed a covalent adduct with VPP, a process that was inhibited by the presence of vancomycin (Fig 8).

**Fig 7.**
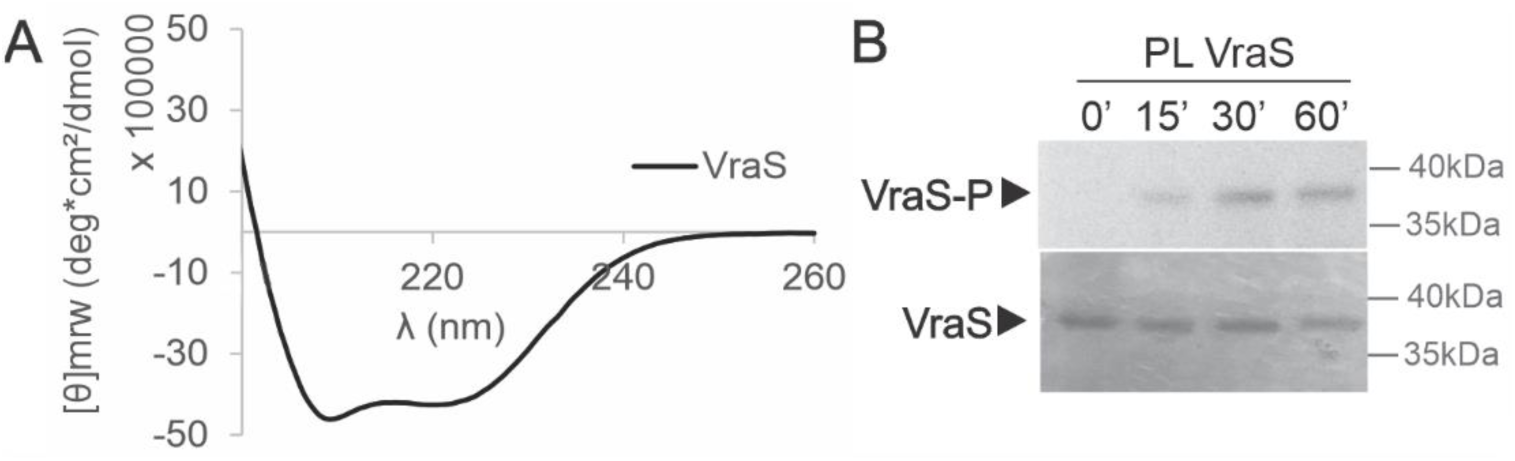
Structural and functional characterization of recombinant VraS. **A.** Circular dichroism spectrum of purified VraS-H6x protein (10 µM) in DDM micelles. **B.** Autophosphorylation of purified VraS-H6x after reconstitution into liposomes. VraS-H6x proteoliposomes were incubated with 0.16 µCi/µl of [^32^P]-γ-ATP (250 µCi, 10 µM) at 37 °C for 0, 15, 30 and 60 min. The reaction was stopped by the addition of SB 5X. Proteins from each sample were resolved by 12% SDS-PAGE (2.5 µg of protein in each well), visualized by autoradiography using a Typhoon FLA7000 (GE) (top) and then stained with Coomassie Brilliant Blue (bottom).

**Fig 8.**
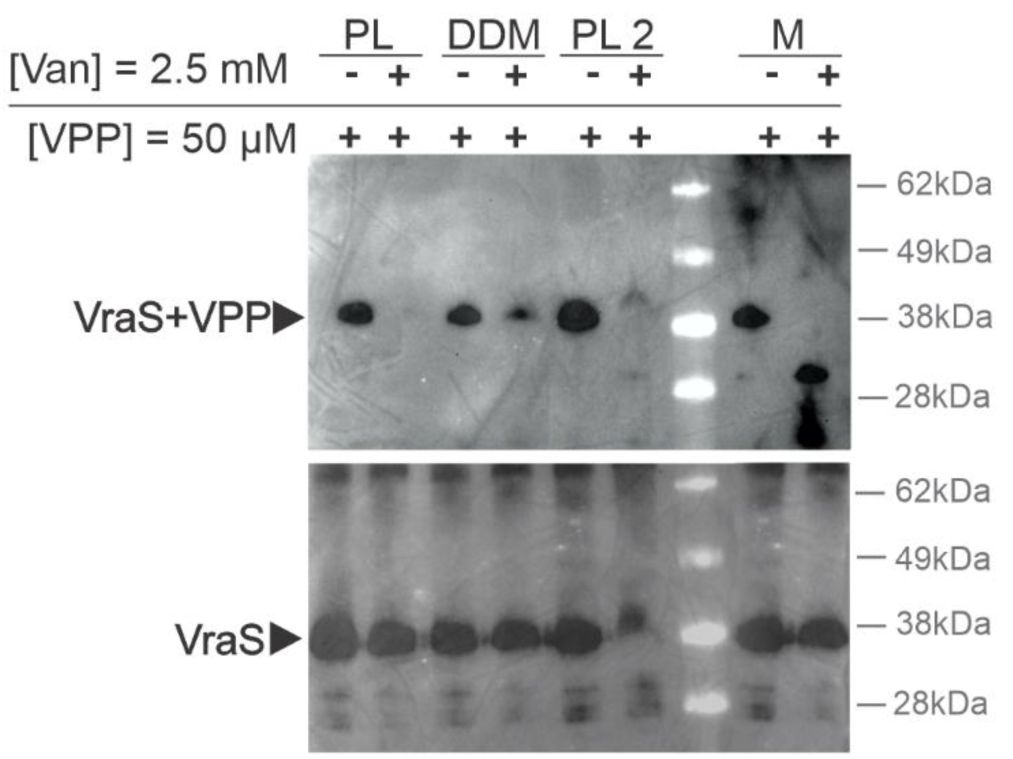
Interaction of purified VraS with VPP in detergent micelles and proteoliposomes. Fractions of VraS proteoliposomes (two different preparations, PL and PL2), purified protein in DDM micelles (DDM), and membrane vesicles over-expressing VraS (M) at a concentration of 0.5 mg/ml of total protein, were pre-incubated with water or 2.5 mM vancomycin for 1 h, then 50 µM of VPP was added and incubated for 1 h. Samples were irradiated at 365 nm for 2 h at 4 °C. Proteins from each sample were resolved by 4-12% NuPAGE (2.5 µg protein in each well) and analyzed by Western blot. Biotin in VPP was detected with Streptavidin-HRP (1/5,000; top), and the His-tag in VraS.H6x was detected with an anti-His-Tag-HRP antibody (1/50,000; bottom). Sequential chemiluminescent detection was performed on the same membrane after HRP inactivation by H_2_O_2_ treatment (1^st^, Streptavidin-HRP; 2^nd^, anti-His-Tag-HRP).

The binding of VPP to VraS was assessed using active protein in proteoliposomes. VraS proteoliposomes were incubated with increasing concentrations of the VPP photoprobe, and after irradiation, the fraction of VPP-bound VraS was determined via Western blot analysis. The results revealed that VPP binding to VraS was concentration-dependent and saturable, with an apparent *K*_D_ of 42 ± 8 μM (Fig 9.A and B). A competition assay with vancomycin further demonstrated the specificity of the interaction, yielding an IC_50_ of 1.6 ± 0.3 mM (Fig 9.C and D).

**Fig 9.**
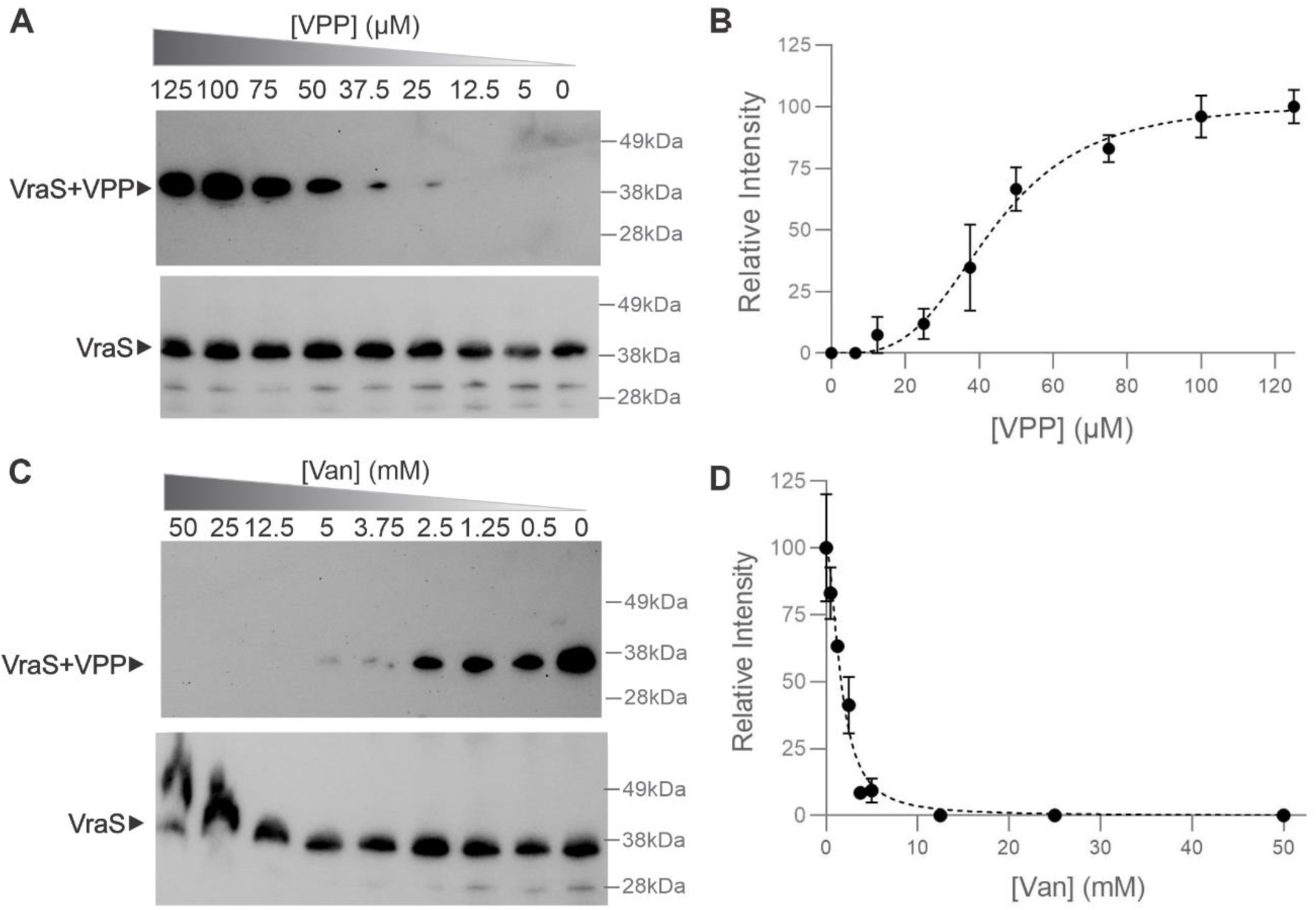
Interaction of VraS with the vancomycin-derived photoprobe and vancomycin competition assays. **A.** VraS proteoliposomes were incubated for 1 h with increasing concentrations of VPP and then irradiated at 365 nm for 2 h at 4 °C. Proteins from each sample were resolved by 4-12% NuPAGE (2.5 µg protein in each well) and analyzed by Western blot. **B.** The protein bands were quantified using the Gel Analyzer software; the intensity of the VPP-bound VraS-H6x protein (evidenced through interaction with Streptavidin-HRP) was relativized to the total amount of VraS-H6x, the band detected in the Western blots revealed with the His-tag specific antibody. The normalized relative signal intensity (Relative Intensity; the fraction of VPP-bound VraS-H6x) was plotted as a function of total VPP concentration. The results represent the means of two experiments, and the error bars correspond to the standard deviations (SD). Experimental points (fraction of VPP-bound VraS-H6x) were fitted to the Hill equation using the GraphPad program, and an apparent *K*_D_ of 42 ± 8 µM (B_max_ = 99 ± 1; h = 4 ± 1) was determined. **C.** VraS proteoliposomes were pre-incubated for 1 h with increasing concentrations of vancomycin; then 50 µM VPP was added, incubated for 1 h, and the samples were irradiated at 365 nm for 2 h at 4 °C. **D.** Quantification of the bands in (C), as described in (B). The normalized relative signal intensity (or Relative Intensity; the fraction of VPP-bound VraS-H6x) was plotted as a function of vancomycin concentration. The results represent the means of two experiments, and the error bars correspond to the standard deviations (SD). Points were fitted to a variable slope inhibitor concentration vs. response curve, and an apparent IC_50_ of 1.6 ± 0.3 mM (Hill Slope =-2 ± 0.5) was determined. In A and C, the biotin moiety in VPP was detected with Streptavidin-HRP (1/5,000; top), and the C-terminal H6x tail of recombinant VraS was detected with a His-Tag specific antibody conjugated to HRP (1/50,000; bottom). Sequential chemiluminescent detection was performed on the same membrane after HRP inactivation by H_2_O_2_ treatment. The duplicate Western blots of the assays shown in (A) and (C) are shown in S5 Fig.

We hypothesize that vancomycin, other glycopeptides, and β-lactams most likely interact with the extracellular loop of VraS or with residues of the TM α-helices located close to the outer leaflet of the membrane. DeepTMHMM [54] was used to predict transmembrane α-helices in VraS. Two transmembrane α-helices were predicted in VraS, spanning residues 8-25 and 45-67 (Fig 1.B and S1 Fig). MembraneFold [55] was used to combine the prediction made with DeepTMHMM with the VraS structure prediction for the monomer deposited in the AlphaFold Protein Structure Database (A0A1L6C0A3; https://alphafold.ebi.ac.uk/; Fig 1.C). Given our experimental evidence for dimer formation, a model of the VraS dimer was generated with AlphaFold3 [56]. In the predicted dimer, the VraS monomers would adopt an antiparallel orientation of the N-terminal regions (Fig 10), with a small interaction surface defined between the short extracellular loops of each monomer (residues 26-44). Residues 26-44 are predicted to connect the two transmembrane α-helices. The C-terminal cytoplasmic core of VraS (amino acids 68 to 347) contains the dimerization domain, which houses the conserved His^156^ residue, and the ATP-binding domain. Through a multiple sequence alignment of the *S. aureus* VraS sequence and a panel of bacterial histidine kinases, the ARELH^156^DSVSQ region was identified, similar to the H-box region of type-III HKs [57].

**Fig 10.**
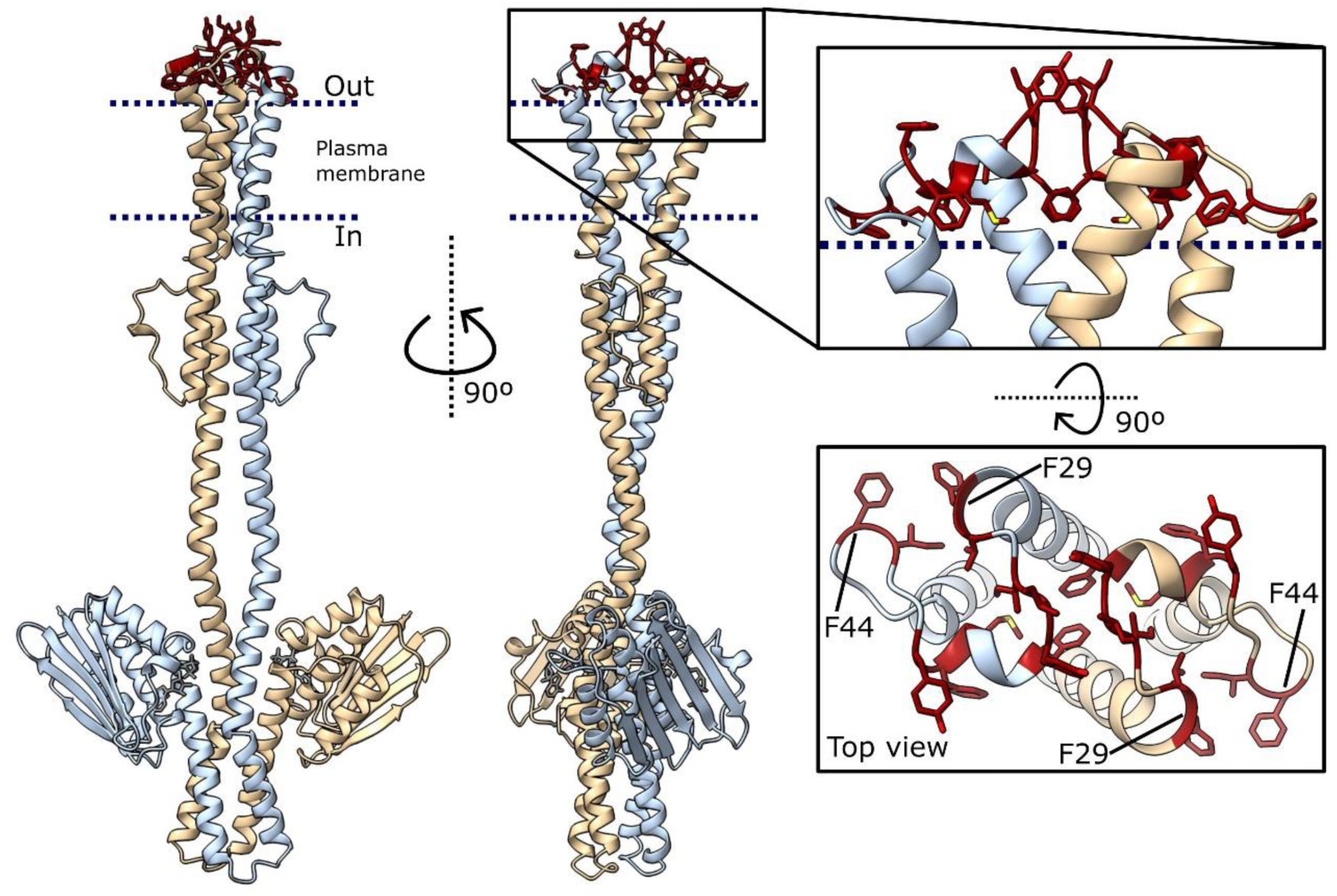
Model of the VraS dimer. AlphaFold3 predicted VraS dimer (one monomer colored yellow and the other one light blue); highlighted in red: the hydrophobic residues in the predicted extracellular loop (residues 26-44); F29 and F44 of each monomer are labelled to show the antiparallel interaction of the monomers.

VraS overexpressing membranes were incubated with VPP (or DMSO as control) to further confirm the interaction and attempt characterization of the vancomycin binding site and subsequently irradiated at 365 nm to activate the benzophenone. Afterward, VraS was solubilized from the membranes and purified using two sequential affinity chromatography steps. The first purification step, using a Ni Sepharose® column, allowed the purification of recombinant VraS through its C-terminal H6x-tag. The eluted protein retained the biotin label, indicating that at least a fraction of the purified VraS protein formed the VraS-VPP adduct (Fig 11.A). Although the active biradical −generated upon irradiation of the benzophenone-bearing vancomycin photoprobe− is generated after irradiation, covalent binding to a biomolecule occurs with low efficiency [58]. Hence, the VraS protein that was affinity-purified using the Ni Sepharose® column (hereafter referred to as VraS+VPP) was most likely a mixture of VraS and of the VraS-VPP adduct, where the latter could be the minority fraction. Intact mass spectra of VraS presented a single peak (S6 Fig), corresponding to a mass of 40,898 Da, in agreement with the theoretical molecular mass for the VraS-H6x protein with the two predicted N-terminal transmembrane helices (40,867.37 Da). In the VraS+VPP sample, no second peak was identified that could correspond to the VraS-VPP adduct (expected mass *ca*. 42.8 kDa; VPP is 1923 Da [40]), possibly due to the low abundance of the VraS-VPP adduct. The VraS-VPP adduct was enriched by subjecting the VraS+VPP sample to a second affinity chromatography step using a streptavidin resin. The VPP-associated biotin label was only identified in the VraS fractions retained and then eluted from the streptavidin resin (Fig 11.B). The amount of protein recovered did not allow for the determination of the intact mass of the adduct. Hence, we dedicated efforts to attempt the identification of the VraS peptide covalently bound to VPP.

**Fig 11.**
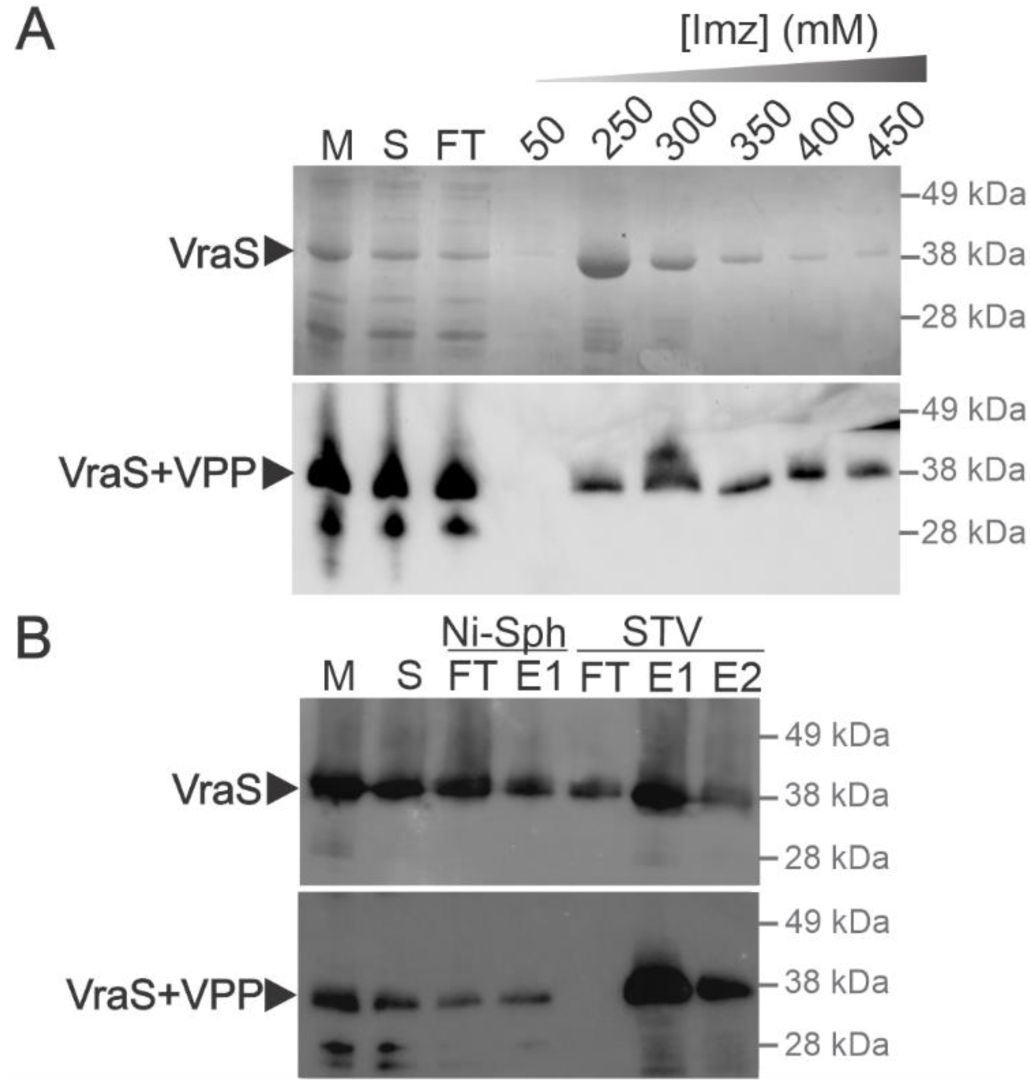
VraS pull-down with the vancomycin-derived photo-affinity probe. **A.** VraS-H6x expressing membrane vesicle preparations were incubated with 50 μM VPP or DMSO (control; same volume used to add the DMSO-dissolved photoprobe) for 1 h and subsequently irradiated at 365 nm for 2 h at 4 °C. Afterwards, VraS-H6x was solubilized with DDM and purified using a Ni Sepharose® 6 Fast Flow resin. The gel shows fractions from the VraS+VPP protein purification process, resolved in a 4-12% NuPAGE gel stained and visualized with Coomassie Brilliant Blue (top) and by Western blot (detection with Streptavidin-HRP, 1/5,000; bottom). M, membranes after incubation and irradiation; S, soluble fraction; FT, Flow-through; [Imz], elution fractions with increasing concentrations of Imidazole (50-450 mM). **B.** Purification of the VraS-VPP adduct using a Streptavidin resin. The purified VraS+VPP fraction (from the Ni Sepharose® 6 Fast Flow column) was incubated with a Streptavidin resin overnight, washed with 25 mM Tris-HCl, 300 mM NaCl (pH 8.5) and eluted after incubation for 30 minutes with 25 mM Tris-HCl, 300 mM NaCl (pH 8.5), 4 mg/ml biotin, 0.1% RapiGest™ SF and subsequent heating at 50 °C for 15 min. Protein fractions from different steps of the purification procedure were resolved in a 4-12% NuPAGE gel and visualized by Western blot. Top: detection with a His-Tag specific antibody HRP-conjugated (α-H6X, 1/50,000). Bottom: detection with Streptavidin-HRP (STV-HRP, 1/5,000). Sequential chemiluminescent detection was performed by inactivation of HRP by treatment with H_2_O_2_). M, membranes after incubation and irradiation; S, soluble fraction; FT, Flow-through, E1-E2, elutions; Ni-Sph, protein fractions from the purification using a Ni Sepharose® 6 Fast Flow resin; STV, protein fractions from the purification using a Streptavidin resin.

We analyzed the VraS peptides generated by in-gel and in-solution digestion assays with trypsin and chymotrypsin. The tryptic and chymotryptic peptides were separated by reversed-phase LC and characterized by MS/MS. The analysis of the peptides generated after in-solution digestion of the VraS and VraS+VPP samples with chymotrypsin and trypsin yielded a sequence coverage of approximately 60-70% (S7-S8 Fig). Sequence coverage comprised amino acid residues 70 to 353; peptides corresponding to the N-terminus of VraS-H6x could not be detected. Sequence coverage was similar for VraS and VraS+VPP. The comparison of the peptides of the VraS and VraS+VPP samples did not allow the identification of any VPP-modified peptide. Although the sequence coverage upon trypsin and chymotrypsin in-solution digestion was very similar, analysis of the streptavidin-purified VraS-VPP fraction was carried out after digestion with chymotrypsin since it presented a greater number of theoretical cut sites at the N-terminal end of the protein (S7 Fig). Sequence coverage after digestion of streptavidin-purified VraS-VPP with chymotrypsin in solution was 64% (S9 Fig), but neither peptides from the N-terminal region nor VPP-labelled peptides were identified.

### Evaluation of the interaction of vancomycin and ampicillin with VraS using Saturation Transfer Difference NMR spectroscopy

Saturation transfer difference (STD) NMR spectroscopy experiments were conducted to confirm the interaction of vancomycin with VraS in solution. STD spectra were calculated from the difference between a reference spectrum (saturating off-resonance at-40 ppm) and a spectrum saturating the methyl groups of the protein at 0.75 ppm (on-resonance). If the protein interacts with a ligand, saturation is transferred from the protein protons to the protons of the ligand due to proximity in space through the Nuclear Overhauser effect. The difference spectrum obtained between the on-resonance and the off-resonance spectra evidences the effect of the saturation transferred from the protein to the ligand. As a protein-ligand interaction is inferred through the evaluation of NMR resonances of the free ligand, the STD assay applies to the assessment of the interaction of ligands with large proteins and proteins in detergent micelles, as is the case of VraS, which is *ca.* 40 kDa in the monomeric form but is present as a dimer in DDM micelles (S4 Fig). The STD spectra obtained from a sample of vancomycin incubated with VraS showed that the aromatic protons of vancomycin were perturbed upon saturation of the methyl group protons of the protein, which confirmed the interaction of the glycopeptide with VraS in solution (Fig 12). The signals of vancomycin in the aromatic region of the spectra were absent in the difference spectra of the antibiotic separately (Fig 12 and S10 Fig) and the protein (S10 Fig). The ^1^H NMR spectra and complete STD spectra are shown in S10 Fig.

**Fig 12.**
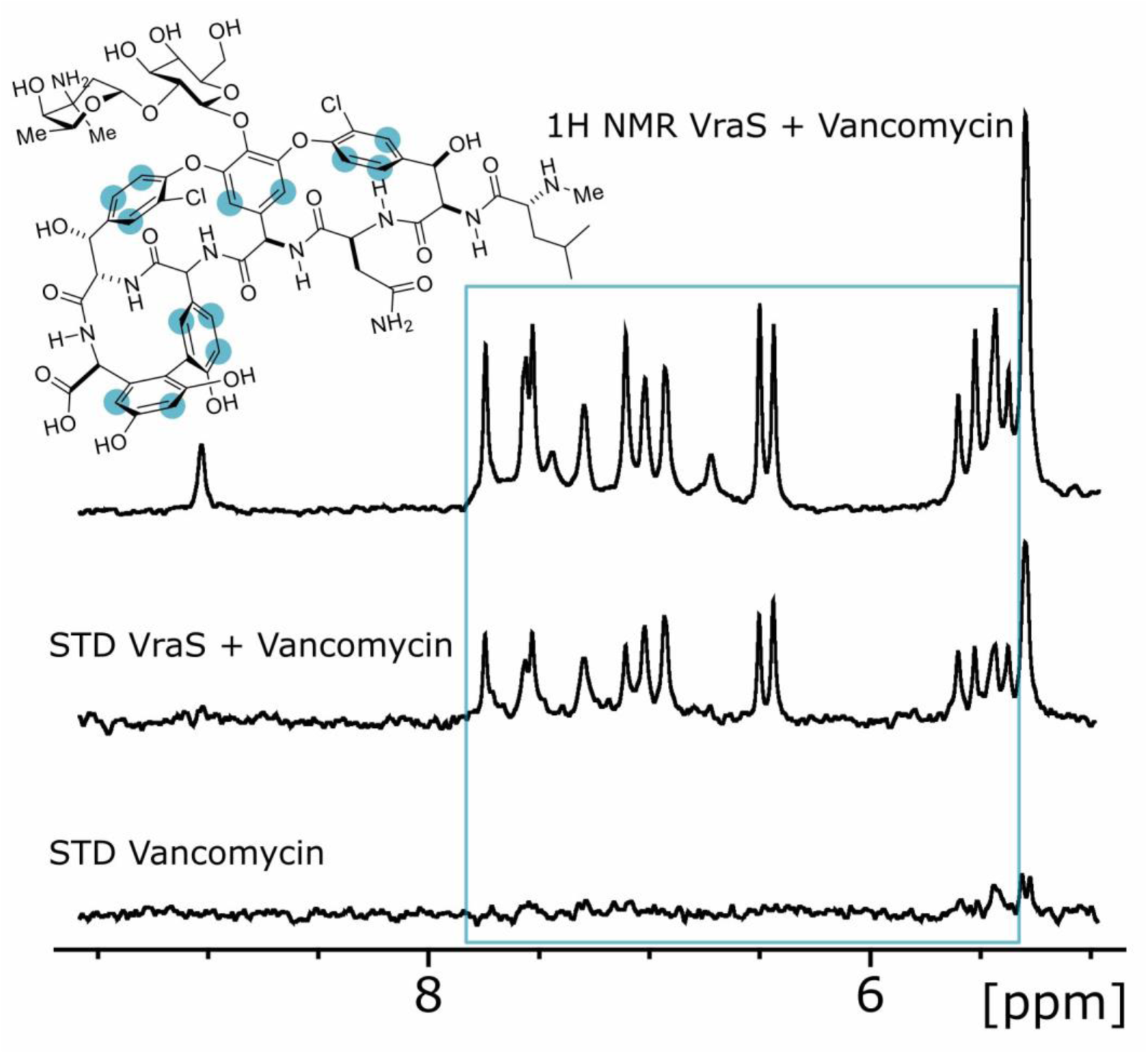
Saturation Transfer Difference NMR assays confirm binding of vancomycin to VraS. ^1^H NMR spectrum of 20 µM full-length VraS in the presence of 400 µM Vancomycin (top), and Saturation Transfer Difference NMR spectra, saturating at 0.75 ppm, of 20 µM full-length VraS with 400 µM vancomycin (middle) and of 400 µM vancomycin (bottom). The peaks inside the blue rectangle, in the ^1^H NMR spectrum of the mixture of VraS with vancomycin, correspond to the aromatic protons of vancomycin (highlighted with a light-blue circle in the structure), and they are only detected in the STD experiment of the mixture, but not in the STD spectrum of vancomycin alone. Buffer was 20 mM sodium phosphate pH 7.4, 280 mM NaCl, 6 mM *KCl,* 0.01% DDM, 10% deuterium oxide.

It has been reported and confirmed here (Fig 2) that β-lactam antibiotics also activate the VraTSR system. Hence, we performed the saturation transfer difference experiments for ampicillin in the presence of VraS. When saturating at 0.75 ppm, the difference spectrum demonstrated the interaction of ampicillin with VraS (Fig 13). The STD spectrum evidenced a change in the signal intensity from the protons of the aromatic ring and the methine proton on carbon 16, but there is no change in the intensity of the signal corresponding to the proton bound to carbon 3. This suggested that ampicillin interacted with VraS preferentially through the substituent bound to carbon 6 of the β-lactam ring. The ampicillin and ampicillin-VraS ^1^H NMR spectra and complete STD spectra are shown in S11 Fig. An STD experiment of kanamycin, one of the antibiotics that did not activate the VraTRS system, showed no evidence of interaction with VraS (S12 Fig).

**Fig 13.**
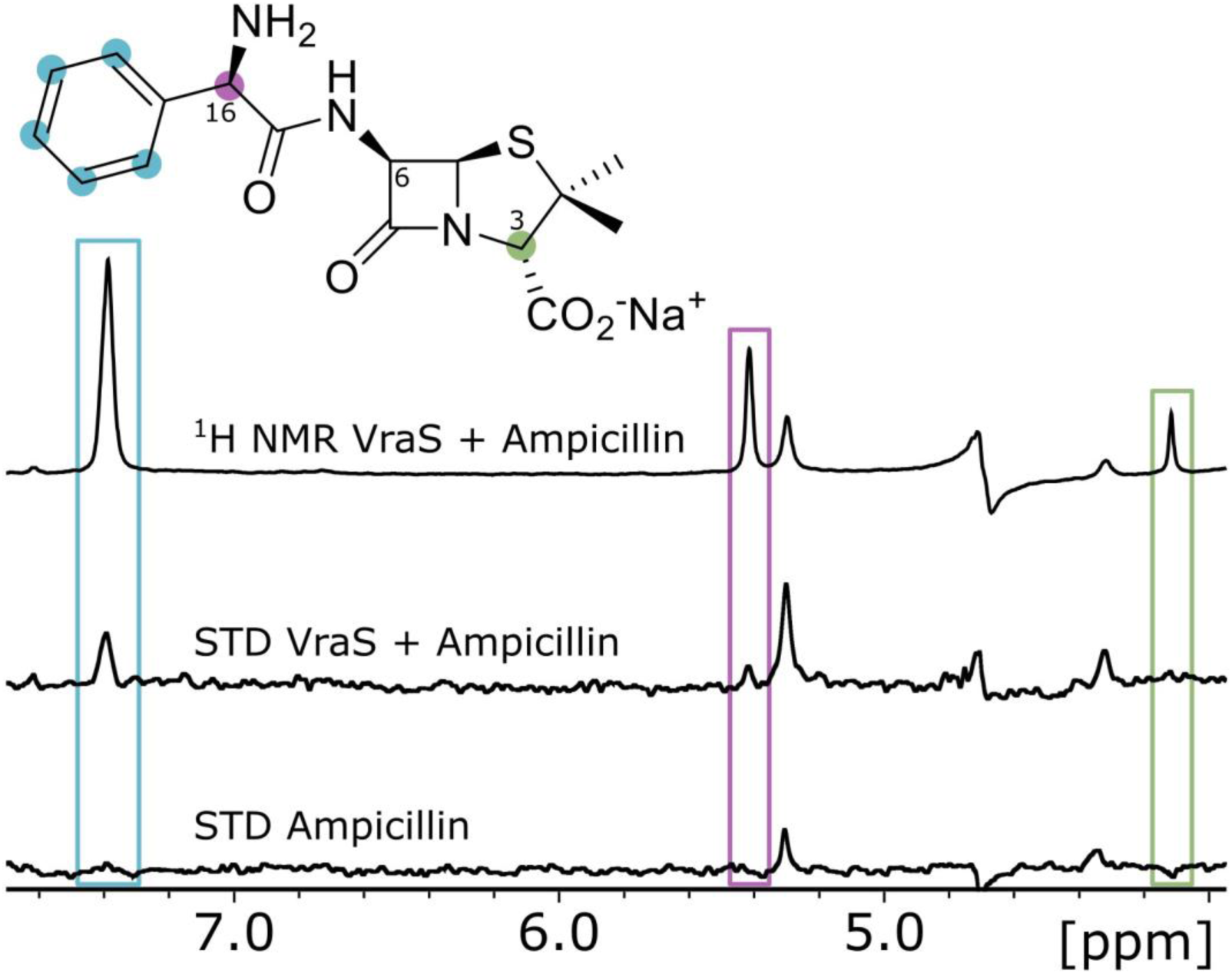
Saturation transfer difference NMR determines binding of ampicillin to VraS. ^1^H NMR spectrum of 20 µM full-length VraS in the presence of 400 µM ampicillin (top), and Saturation Transfer Difference NMR spectra, saturating at 0.75 ppm, of 20 µM full-length VraS with 400 µM ampicillin (middle) and of 400 µM ampicillin (bottom). The peaks inside the blue and purple rectangles in the ^1^H NMR spectrum of the mixture of VraS with ampicillin correspond to the aromatic protons (blue circle in the structure) and the methine proton on carbon 16 of the β-lactam (purple circle). They are only detected in the STD experiment of the mixture but not in the STD spectrum of vancomycin alone. These STD assays show the substituent on carbon 6 of ampicillin interacted with VraS. Instead, the green rectangle indicates the peak corresponding to the methine proton on carbon 3 of ampicillin, which was not detected in the STD of the VraS-ampicillin mixture, indicating that this part of the fused ring did not interact with VraS. Buffer was 20 mM sodium phosphate pH 7.4, 280 mM NaCl, 6 mM *KCl,* 0.01% DDM, 10% deuterium oxide.

## Discussion

Two-component systems (TCS) allow the adaptation of bacteria to changing environments [59–61]. The TCSs involved in activating antibiotic resistance mechanisms are of particular interest to address recalcitrant bacterial infections. Inhibition of the sensor histidine kinases or the phosphorylated response regulators could restore the activity of antibiotics that have lost their efficacy due to resistance [60]. Most histidine kinases are transmembrane receptors, with an N-terminal signal sequence serving as a first transmembrane helix, a periplasmic or extracellular sensor domain, and a second transmembrane helix. A wide variety of domains in the periplasm or extracellular space sense directly diverse signals (*e.g.,* small ligand nutrients such as amino acids and carboxylic acids) and regulate histidine kinase activity [61]. Examples are the PAS domains of CitA (citrate ligand) [62], the tandem PAS domain of KinD (pyruvate ligand) [63], the PAS domain of the *Salmonella typhimurium* PhoQ histidine kinase (which detects small antimicrobial peptides and Mg^2+^ ions) [64–66] or the all α-helical domain of NarX (nitrate ligand) [67]. The NreB kinase detects oxygen concentration directly by utilizing a cytoplasmic PAS domain with an [4Fe–4S]^2+^ cluster [68]. In contrast, some membrane-bound histidine kinases have two or more transmembrane segments connected by intra-or extracellular loops of a few amino acids. As they lack any additional domain that would allow signal detection in the periplasmic or extracellular space, it has been proposed that these proteins sense their stimulus either directly inside or at the surface of the membrane [69]. Examples of these intramembrane-sensing histidine kinases are DesK and LiaS of *Bacillus subtilis*. DesK is suggested to detect membrane thickness as an indicator of membrane viscosity, triggering lipid desaturation in response to disturbances. In DesK, it has been proposed that the transmembrane domain senses the changes in membrane thickness and transmits the information to the downstream catalytic domains [70, 71]. LiaS and its homologs (including VraS) are involved in sensing cell envelope stress [70, 72]. The LiaRS system is activated by antibiotics that interfere with the Lipid II cycle of cell-wall peptidoglycan biosynthesis [73] and is considered a three-component system as a third protein, LiaF, acts as a potent inhibitor of LiaR-dependent gene expression in the absence of stimuli [26]. In contrast, VraT is required for VraR-dependent gene expression in the homologous VraTSR system [30, 33]. Yet, other systems use cytoplasmic-sensing histidine kinases (either membrane-anchored or soluble) that detect cellular or diffusible signals reporting the metabolic or developmental state of the cell [74]. Regarding the membrane-bound sensor/transducer kinases of TCS, many display dual activity as, apart from catalyzing phosphorylation of the response regulator, they also catalyze its de-phosphorylation [32, 61, 70, 71].

An intriguing characteristic of the VraTSR system is that it is activated by different classes of cell-wall active antibiotics. Using a GFP-based reporter strain, we corroborated that β-lactam antibiotics and glycopeptides activate the VraTSR system. Regarding the latter class of antibacterials, here we showed that the glycopeptides corbomycin [45], complestatin [45], A47934 [75], and rimomycin A [46, 47] activate the VraTSR system.

Most glycopeptides bind the D-Ala-D-Ala terminal dipeptide of the pentapeptide in Lipid II and glycan strands, inhibiting the transglycosylation and transpeptidation reactions catalyzed by PBPs. However, the antibacterial activity of the Type V glycopeptides corbomycin, complestatin, and rimomycin A results from indirectly inhibiting autolysins by binding peptidoglycan [45–47]. Our results with corbomycin and complestatin agree with recent findings that the VraTSR system is the only TCS whose absence affects the susceptibility of *S. aureus* to these antibiotics [76]. Our assays with the reporter strain also showed that the phosphoglycolipid moenomycin A induces the VraTSR system. Moenomycin A inhibits bacterial cell wall synthesis by binding to the transglycosylases responsible for peptidoglycan carbohydrate chain formation [50–52]. Hence, our results expand the panel of structures of the cell-wall active antibiotics that activate the VraTSR system.

The VraTSR system responds to many antibiotics, whose structures and mechanisms of action are highly variable. For this reason, it has been postulated that the stimulus that triggers resistance would not be antibiotic recognition but rather an intermediate of cell wall synthesis generated by their action [14, 30, 77]: a compound that accumulates because of inhibition of cell-wall synthesis while cell wall recycling continues unperturbed. In this scenario, peptidoglycan fragments and/or intermediates of cell-wall synthesis were proposed as candidate inducers/signals. Here, we demonstrated that compounds that inhibit the autolysins involved in cell wall recycling also activate the VraTSR system and that peptidoglycan fragments generated by the action of lysostaphin, mutanolysin, or the combination of both did not activate the system nor potentiate the action of β-lactams or glycopeptides. While this study probed the effects of glycan chains and peptide stems produced by a muramidase, it was not comprehensive as other peptidoglycan hydrolases, including lytic transglycosylases and glucosaminidases, play roles in peptidoglycan homeostasis of *S. aureus.* Therefore, other peptidoglycan fragments will be explored in future work, including those that are membrane-associated. The structural diversity of VraTSR inducers could reflect the involvement of different proteins in signal detection, and both VraS and VraT, membrane-bound proteins with regions exposed to the extracellular space, could sense antibiotics. VraS has the structural characteristics of an intramembrane-sensing histidine kinase and could be activated in response to a stimulus associated with or occurring directly at the membrane interface, as it has been proposed for intramembrane-sensing histidine kinases lacking a periplasmic domain [32, 69]. Alternatively, VraS activation could be indirect, dependent on protein-protein interactions with VraT [30]. VraS dimerization and/or VraS-VraT oligomerization could lead to ligand binding pockets in protein-protein interfaces [78]. The binding of a ligand can shift the equilibrium between monomers and multimers [79] and modulate activity. Contrary to prevailing assumptions, our findings reveal VraS as a direct receptor for vancomycin and ampicillin, fundamentally reshaping our understanding of antibiotic sensing in the VraTSR system. Although no covalent adducts were formed with VraT, our results do not exclude its potential role in antibiotic detection or signal transduction. VraT is essential for manifesting resistance through a mechanism still to be elucidated. It is conceivable that the VraS-antibiotic complex recruits VraT, a hypothesis that warrants further investigation to unravel the exact activation mechanism of the VraTSR system.

Vancomycin is a very large molecule that cannot cross the plasma membrane, so it cannot enter the cellular cytoplasm of the bacteria. Therefore, biologically, the interaction with the antibiotic could only occur through the extracellular loop, comprising amino acids 26 to 44, and/or through the residues of the transmembrane α-helices inserted in the extracellular leaflet of the membrane. Our efforts to identify the VPP-crosslinked peptide have so far been unsuccessful. However, STD NMR assays with vancomycin and ampicillin revealed interactions between the aromatic rings of these antibiotics and VraS, providing an initial clue to a shared structural feature among the diverse antibiotics that activate VraTSR. The extracellular loop of VraS contains several hydrophobic groups (highlighted in red in Fig 10; the hydrophobic surface is shown in S4 Fig), suggesting the possibility of hydrophobic interactions between the antibiotics and this loop. Further experiments will be necessary to confirm the precise binding site.

By identifying VraS as a receptor for glycopeptides and β-lactams, this study unveils a pivotal mechanism in antibiotic sensing and opens new avenues for combating antibiotic resistance. The demonstrated direct interaction of vancomycin and ampicillin with VraS, supported by STD and competition assays using both the full-length protein and the active protein reconstituted in liposomes, paves the way for the development of novel therapeutics. These could include compounds that compete with glycopeptides and β-lactams for binding or inhibitors that target VraS autophosphorylation.

The ability of diverse cell wall-active antibiotics to activate VraTSR highlights the complexity of its regulatory mechanisms, suggesting the presence of multiple activation pathways. While our findings establish the feasibility of direct activation, they also point to a tantalizing horizon of undiscovered signals—hidden cues that could revolutionize our understanding of bacterial stress responses and inspire innovative antimicrobial strategies.

## Experimental procedures

### Strains, plasmids, and reagents

All chemicals were of the highest quality available and purchased from Sigma-Aldrich, BioRad, Thermo Fisher Scientific, Abcam, and Promega. *Escherichia coli* DH5α (laboratory stock; Table 1) cells were used for transformation with ligation mixtures for cloning and transformation with plasmids for storage as frozen stocks. *E. coli* BL21 Star^TM^ (DE3) cells (laboratory stock; Table 1) were employed for protein production from genes cloned in pET24a(+). *Staphylococcus aureus* RN4220 cells (laboratory stock; Table 1) were used for transformation with pCN52-derived plasmids. Luria-Bertani (LB) broth (Difco) was used as growth medium for all bacterial strains. VPP was synthesized using a method already reported (40).

**Table 1.** Bacterial strains.

| Name | Characteristic | Reference |
| --- | --- | --- |
| *E. coli* DH5α | *F*- *endA1 glnV44 thi-1 recA1 relA1 gyrA96 deoR nupG*<br>Φ80*dlacZ*ΔM15 $\Delta$(*lacZYA-argF*)U169, *hsdR17*(*rK*-*mK*+), λ- | |
| *E. coli* BL21 Star<sup>TM</sup> (DE3) | *F*- *ompT hsdSB (rB-mB-) gal dcm rne131* (DE3) |  |
| *S. aureus* RN4220 | Restriction-deficient derivative of NCTC8325-4 | [80] |

### Genes and oligonucleotides

The *vraS* and *vraT* synthetic genes, optimized for expression in *E. coli*, were purchased from Genscript (Table 2). The *vraS* and *vraT* genes and the operator region of the VraTSR system from *S. aureus* USA300-TCH1516 were synthesized without restriction sites, and the sites required for cloning were incorporated in the oligonucleotides used in the PCRs. The *vraT* gene was synthesized to include a Twin-Strep-Tag in the C-terminal of the encoded protein. The oligonucleotides (Table 3) were purchased from Genbiotech.

**Table 2.**
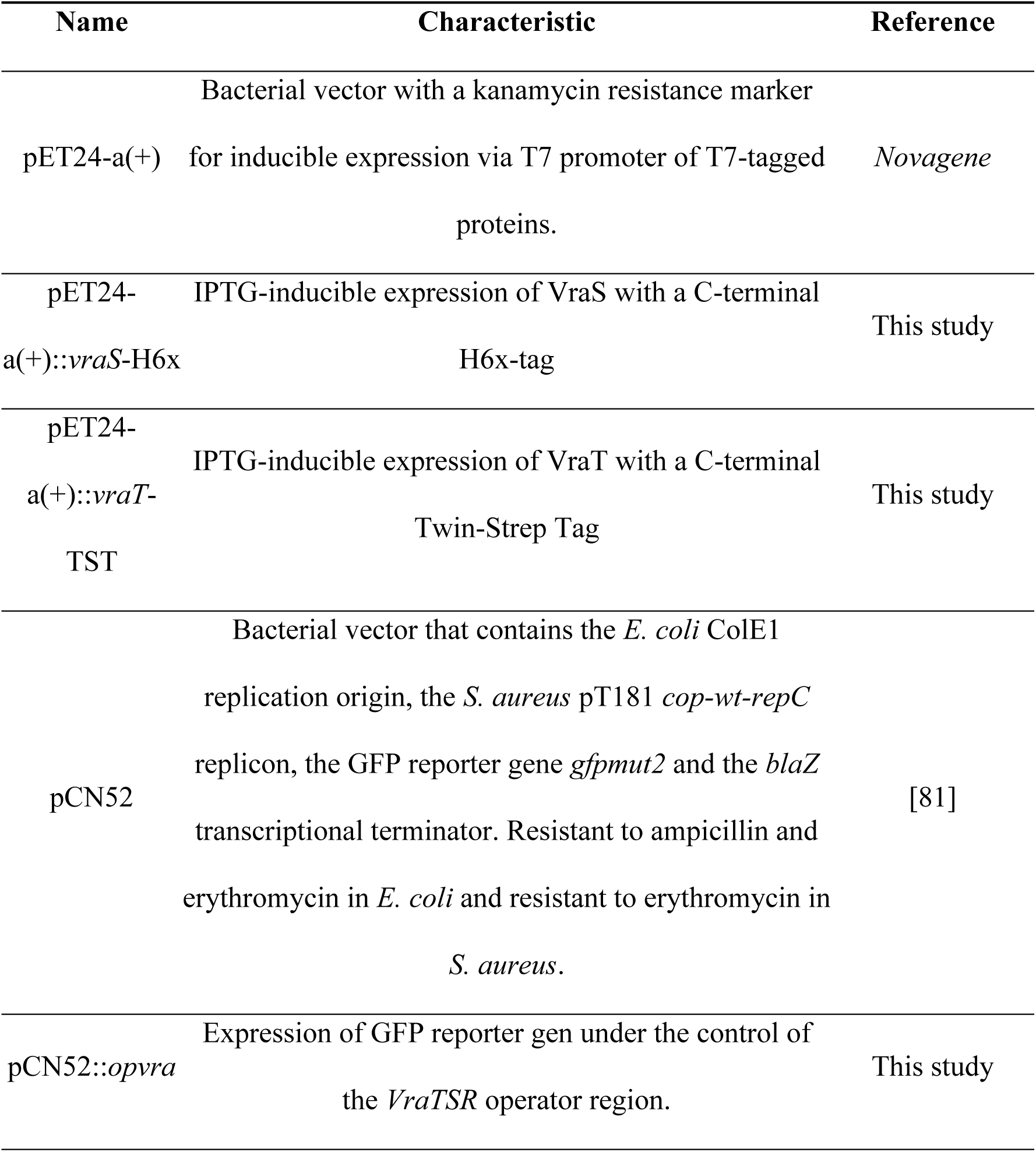
Plasmids.

**Table 3.**
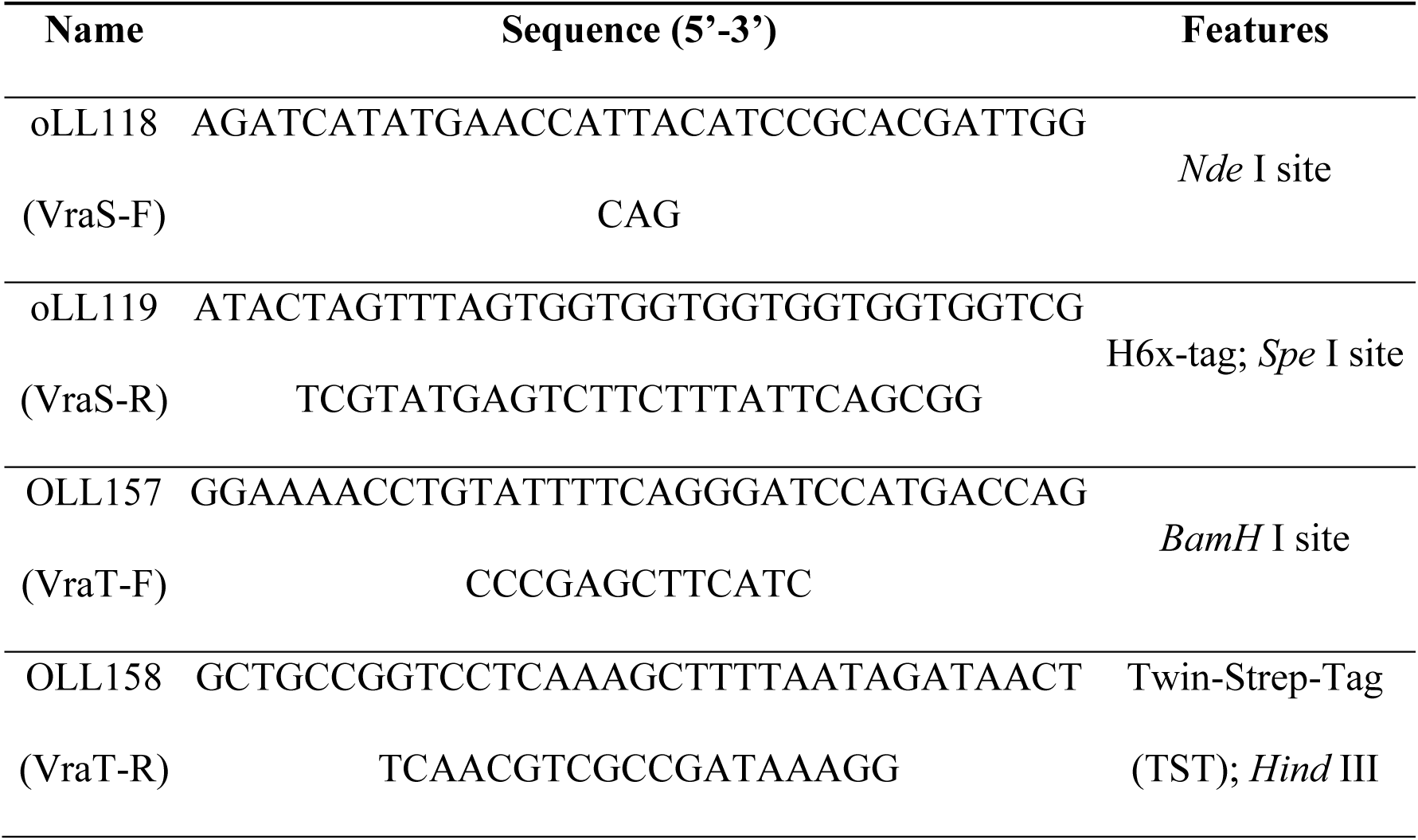
Oligonucleotides used as primers in PCR reactions.

### DNA Techniques and Cloning Procedure

DNA preparation and related techniques were performed according to standard protocol (Sambrook J, Fritsch EF, & Maniatis T (1989) Molecular Cloning: A Laboratory Manual (Cold Spring Harbor, NY) 2nd Ed.). Plasmid DNA was isolated using the Wizard Plus SV Minipreps kit (Promega). Purification of PCR products and plasmids from agarose gels was carried out using the Wizard® SV Gel and PCR Clean-Up System kit (Promega). All restriction enzymes were purchased from Promega; T4 DNA ligase (Promega) and Platinum Pfx DNA Polymerase (Thermo Fisher Scientific) were used according to the instructions in the corresponding user manual. The constructs were verified through sequencing at the University of Maine DNA Sequencing Facility (https://umaine.edu/dnaseq/).

### Construction of *Staphylococcus aureus* VraTSR reporter strain

To construct the *vraTSR* promoter fusion, a 259-bp synthetic DNA fragment was acquired from GenScript, cloned in pUC57 plasmid. This synthetic DNA fragment contained the 247-bp DNA fragment of the *vra*TSR operator region (including the-121 to +26 region [43] and extending up the last nucleotide before the ATG of the *vraU* gene), flanked by *Sph* I and *Eco*R I sites. The 259-bp fragment was purified after digestion of the pUC57::*opvra* plasmid with *Sph* I and *Eco*R I, and was ligated upstream of the *gfp* gen in plasmid pCN52. *S. aureus* RN4220 was transformed by electroporation with pCN52::*opvra* (to generate the reporter strain) or pCN52 (as a control for basal level of fluorescence of the cells). Electroporation was carried out with a Gene Pulser Xcell Electroporation System (Biorad), operated at 200 Ω resistance, 2.5 kV voltage and 25 µF capacitance using 0.2 cm gap cuvette (Gene Pulser/MicroPulser Electroporation Cuvettes, Biorad). Cells were recovered in LB for 1h at 37 °C, 200 rpm. Transformants were selected in LB-erythromycin-agar plates.

To monitor activation of VraTSR system, *S. aureus* RN4220 cells transformed with plasmid pCN52::*opvra* or pCN52 (control) were growth at 37 °C, 220 rpm until OD_600nm_ reached 0.3, at which point aliquots of 200 µl of culture were placed in wells of a 96-wells plate and induced with different concentrations of antibiotics (0 to 40 µg/ml), photoprobes (0 to 40 µg/ml) and/or peptidoglycan fragments. Optical density at 600 nm and fluorescence emission of GFP (excitation filter at λ = 485 ± 20 nm and emission filter at λ = 528 ± 20 nm) were simultaneously monitored every 10 min for 2 h in a Synergy Neo2 Hybrid Multi-Mode Microplate Reader (BioTek). The fluorescence intensity (F) was relativized to the number of cells in the culture (OD_600 nm_), after a given time from the addition of the antibiotic/photoprobe/peptidoglycan fragment. To eliminate fluorescence not due to activation of the VraSRT system (auto-fluorescence of *S. aureus* cells), the F/OD of the strain with the empty vector was subtracted, calculating Δ(F/OD) = F/OD _pCN52::*opvra*_ – F/OD _pCN52_. The graphs represent the change in relative fluorescence at each time, Δ(F/OD)_t,_ which corresponds to [(F/OD_pCN52::*opvra*_ – F/OD_pCN52_)_t_ – (F/OD_pCN52::*opvra*_ – F/OD_pCN52_)_t=0h_]. Alternatively, the graphs show the change in relative fluorescence after 2 hours of incubation, Δ(F/OD)_2h_, which corresponds to [(F/OD_pCN52::*opvra*_ – F/OD_pCN52_)_t=2h_ – (F/OD_pCN52::*opvra*_ – F/OD_pCN52_)_t=0h_]. The viability of the cells was checked once the incubation with antibiotics had ended, by means of successive passages in medium without antibiotic and subsequent plating, obtaining a comparable number of viable cells in all cases.

To evaluate induction of the VraTSR system by ampicillin, oxacillin or vancomycin, increasing concentrations of the corresponding antibiotic (0.15, 0.25, 0.5, 1, 2.5, 5 and 10 µg/ml) diluted in water were added. For the evaluation of other antibiotics that act at the cell wall level, as well as antibiotics that act by interfering with other processes, controls were carried out where the cells were supplemented with the same volume of solvent in which the antibiotic is dissolved. It should be noted that the cells that were incubated with daptomycin were also supplemented with CaCl_2_ at a final concentration of 50 µg/ml.

### VraS Topology Prediction and VraS dimer modeling

Transmembrane α-helices in VraS were predicted using DeepTMHMM [54] (https://dtu.biolib.com/DeepTMHMM). MembraneFold [55] (https://ku.biolib.com/MembraneFold/) was used to combine the transmembrane protein topology prediction made with DeepTMHMM with the protein structure prediction for VraS monomer deposited in the AlphaFold Protein Structure Database (A0A1L6C0A3; https://alphafold.ebi.ac.uk/). The VraS dimer was modeled using AlphaFold3 [56].

### Expression of recombinant VraS

The *vraS* gen was amplified by PCR with the *vraS*-F primer, which adds the *Nde* I restriction site, and the *vraS*-R primer, which adds the H6x-Tag and the *Spe* I restriction site (Table 3). The amplification product was digested with the corresponding restriction enzymes and ligated in plasmid pET24-a(+) digested with the same enzymes. CaCl_2_-chemically competent *E. coli* DH5α cells were transformed with the ligation mixture and transformant colonies were selected in LB-kanamycin-agar plates. The construction was checked by sequencing the insert in the purified plasmid. The plasmid pET24-a(+)::*vraS-H6x* allowed isopropyl β-D-1-thiogalactopyranoside (IPTG)-inducible expression of VraS. To express VraS-H6x, *E. coli* BL21 Star^TM^ (DE3) cells harboring the pET24-a(+)::*vraS-H6X* vector were grown overnight in LB broth supplemented with 50 µg/ml kanamycin, at 37 °C, 220 rpm. Fresh medium was inoculated with the overnight culture (dilution of 1/100) and the culture was grown at 37 °C until the OD_600 nm_ reached 0.7, at which point the cultures were rapidly cooled to 20 °C and induced with 50 µM IPTG. VraS expression was induced overnight (16-20 h) at 20 °C, 220 rpm.

### Expression of recombinant VraT

The synthetic gene *vraT_opt* was optimized by GenScript from the sequence encoding the VraT protein from USA300-TCH1516. The optimized gene included an *Nde* I restriction site at 5’ and the coding sequence for the Twin-Strep-Tag (TST) at 3’, followed by a stop codon and the *Spe* I and *Hind* III cleavage sites. The pUC57 vector harboring the synthetic *vraT_opt* gen was digested with the restriction enzymes *Nde* I and *Hind* III and ligated with the expression plasmid pET24-a(+) digested with the same enzymes. CaCl_2_-chemically competent *E. coli* DH5α cells were transformed with the ligation mixture and transformant colonies were selected in LB-kanamycin-agar plates. The construction was checked by sequencing the insert in the purified plasmid. The plasmid pET24-a(+)::*vraT-TST* allowed IPTG-inducible expression of the VraT. *E. coli* BL21 Star^TM^ (DE3) cells harboring pET24-a(+)::*vraT-TST* vector were grown overnight in LB broth supplemented with 50 µg/ml kanamycin, at 37 °C, 220 rpm. Fresh medium was inoculated with the overnight culture (dilution of 1/100) and grown at 37 °C until the OD_600 nm_ reached 0.7, at which point the cultures were rapidly cooled to 20 °C and induced with 500 µM IPTG. VraT expression was induced overnight (16-20 h) at 20 °C, 220 rpm.

### Co-expression of recombinant VraT and VraS

pET24-a(+)::*vraS* was digested with *Xba* I and *Hind* III; the DNA fragment corresponding to RSB-*vraS*-H6x was purified and cloned in pET24-a(+)::vraT-TST previously digested with *Spe* I and *Hind* III, to generate pET24-a(+)::*vraT*-TST-*vraS*-H6x. CaCl_2_-chemically competent *E. coli* DH5α cells were transformed and colonies were selected in LB-kanamycin-agar plates and cloning was confirmed by sequencing the inserts in the purified plasmid. The construct allowed IPTG-inducible co-expression of both proteins, VraS-H6X and VraT-TST. *E. coli* BL21 Star^TM^ (DE3) cells harboring this vector were grown overnight in LB broth supplemented with 50 µg/ml kanamycin, at 37 °C, 220 rpm. Fresh medium was inoculated with the overnight culture (dilution of 1/100) and grown at 37 °C until the OD_600 nm_ reached 0.7, at which point the cultures were rapidly cooled to 20 °C and induced with 50 µM of IPTG and the cells were grown overnight (16-20 h) at 20 °C, 220 rpm.

### Preparation of membrane protein extracts

The cells from 100-ml *E. coli* BL21 Star^TM^ (DE3) cultures expressing the protein of interest were harvested by centrifugation for 15 min at 4,700 *g*, 4 °C in a Sorvall ST 16R refrigerated centrifuge (Thermo Scientific). The cell pellet was then washed with 25 ml of Phosphate Buffer 1 solution (10 mM NaH_2_PO4, 280 mM NaCl, 6 mM KCl, pH 7.4) and the cells were resuspended in 5 ml of Phosphate Buffer 1. The cells were then disrupted by sonication.

For this, the cell culture was supplemented with 2 mM phenylmethylsulfonyl fluoride (PMSF) and 0.01 mM EDTA and subjected to 10 pulses with 20% power in a 600-Watt Sonics GEX-600 sonicator (Sonics & Materials). Each pulse was applied for 15 s, leaving 60 s intervals between pulses, keeping the solution in a water-ice bath. This lysate was centrifuged for 20 min at 12,000 *g* in a refrigerated centrifuge at 4 °C. The supernatant was subjected to ultracentrifugation for 1 h at 150,000 *g* at 4 °C in an OptimaTM L-90K ultracentrifuge (Beckman Coulter), using a Ti90 rotor. The supernatant containing the cytoplasmic and periplasmic protein fraction was separated, while the pellet (containing the fraction of vesicles with membrane proteins) was resuspended in 1 ml of buffer (10 mM HEPES, pH 7.5 or 10 mM sodium phosphate pH 7.4, 155 mM NaCl, 3 mM KCl, depending on the buffer required for the specific assay, as specified below) using a Potter-Elvehjem PTFE pestle and glass tube (SIGMA). The fraction of vesicles and membrane proteins were quantified using the Pierce™ BCA Protein Assay kit (Thermo Scientific) in a microplate by absorbance at 560 nm, according to the supplier’s instructions. Vesicles and membrane protein fractions prepared at McMaster University were performed following the same protocol, using the Model 505 Sonic Dismembrator (Fisherbrand), Sorvall WX100 ultracentrifuge with T-865 rotor (Thermo Fisher Scientific). In this case, the quantifications were performed using the Bradford method (BIORAD) in microplates by absorbance at 595 nm, according to the supplier’s instructions. If the membrane protein extracts were not used right after being obtained, they were fractionated and frozen with liquid nitrogen for storage at - 80 °C. For larger-scale purification, 3.5 l of culture was used, and the protocol was scaled accordingly.

### Evaluation of the interaction of the vancomycin-derived photoprobe with VraS and VraT in membrane protein Extracts

Cultures of *E. coli* BL21 Star^TM^ (DE3) transformed with the empty pET24a(+) plasmid, or with plasmids allowing overexpression of VraS, VraT, and VraS/VraT were grown and protein expression was induced as described above. A membrane protein preparation containing the protein of interest (total protein concentration of 1 mg/ml, in 10 mM HEPES buffer, pH 7.5) was divided into 4 aliquots and 50-µL aliquots were placed in wells in a 96-well microplate. Vancomycin was added to one of these wells (well 1) at a final concentration of 2.5 mM, while an equal volume of water was added to the other three wells and incubated for 1 h at room temperature, protected from light. After that time, the vancomycin-derived photoprobe (VPP) was added at a final concentration of 50 μM to the protein samples in wells 1 and 2, while the Dummy photoprobe was added in a final concentration of 50 μM to the protein sample in well 3 and equal volume of DMSO (solvent in which the photoprobes were dissolved) was added to the sample in the fourth well. The plate was incubated for 1 h at room temperature protected from light and then the plate was placed on ice and irradiated for 2 h with UV-light (λ = 365 nm; Spectronics ENF-240C Handheld UV Lamp). After the irradiation period, 5X protein loading buffer was added to each well and the proteins were resolved by SDS-PAGE followed by Western blot with biotin-specific antibodies to detect biotinylated proteins (including the VPP-bound proteins), with His-tag-specific antibody (HRP-conjugated) to detect VraS-H6x, or with Strep-Tactin-HRP to detect VraT-TST and also VPP-labelled proteins.

### Western Blot Assays

Proteins resolved in 12% Tris-glycine SDS-PAGE or 4-12% NuPAGE gels were transferred to 0.45 µM nitrocellulose membranes (BioRad) using Transference Buffer (25 mM Tris, 192 mM Glycine, 20% Methanol, pH 8.3) in a Trans-Blot system (BioRad). The membranes were blocked overnight at 4°C in 10 ml of Blotto (100 mM Tris-HCl, pH 7.5, 150 mM NaCl, 3% bovine serum albumin, 3% non-fat milk, 0.02% sodium azide,). Incubation of the membrane was performed with the specific antibody for 1 h at RT with gentle agitation: for VraS detection a rabbit His-Tag specific antibody conjugated to HRP was used (Abcam Ab1187; 1/50,000 dilution); for VraT detection, we used Precision Protein Strep-Tactin-HRP Conjugate (BioRad #161-0381, dilution 1/5000); for detection of biotin in VPP, a anti-biotin antibody produced in goat (Sigma-Aldrich B3640, dilution 1/5,000) as a primary antibody and xxx-anti-goat HRP-conjugated antibody (Thermofisher Scientific, dilution 1/5,000) as a secondary antibody. Prior and after the incubation with the antibodies, the membranes were washed 4 times with Tween/Tris-buffered saline (T-TBS; 100 mM Tris-HCl, pH 7.5, 150 mM NaCl, 0.1% Tween 20). The enhanced chemiluminescence system Super-Signal West Pico substrate (Thermofisher Scientific) was used according to the manufacturer’s instructions to detect the immune complexes, together with CL-XPosureTM films (Thermofisher Scientific). The antibodies were diluted in Tween/Tris-buffered saline with 1.2% nonfat dry milk in all cases. The molecular markers used were the PageRuler Prestained Protein Ladder (Thermofisher Scientific) or Blue Plus Protein Ladder (AP Biotech).

### Saturation transfer difference spectroscopy experiments

Saturation transfer difference spectroscopy (STD) experiments were acquired on a Bruker Avance III 700 MHz spectrometer, employing standard Bruker stddiffesgp.3 pulse program. The protein saturation pulse was Sinc1.1000 with 50 ms duration. Saturation time was 2 sec, and relaxation delay was 3 sec. 384 scans and 4 averages were acquired. Off-resonance saturation frequency was-40 ppm and on-resonance saturation frequency was 0.75 ppm. Protein concentration was 20 µM and ligand concentration was 400 µM in a 500 µl final volume. Buffer composition was 10 mM sodium phosphate pH 7.4, 280 mM NaCl, 6 mM KCl, 0.01% DDM, 10% deuterium oxide.

### Purification of VraS from membrane protein extracts

Membrane vesicles expressing VraS were prepared as previously described. VraS-H6x solubilization was performed by incubation of the membrane vesicles with 0.5% n-dodecyl-β-D-maltoside (DDM) in buffer: for circular dichroism spectroscopy, the protein was solubilized in 10 mM sodium phosphate pH 7.4, 155 mM NaCl, 3 mM KCl, 0.5% DDM; for preparation of proteoliposomes the protein was solubilized in 10 mM HEPES buffer, pH 7.5, 0.5% DDM. A 2-ml membrane protein extract (obtained from 2 liters of culture) was diluted 10 times in buffer and then supplemented with the same volume (20 ml) of a 1% (2X) DDM solution in the corresponding buffer (10 mM HEPES solution, pH 7.5 or 10 mM sodium phosphate pH 7.4, 155 mM NaCl, 3 mM KCl −here after referred to as Phosphate Buffer 2−). The mixture was incubated at 4 °C with gentle shaking (top to bottom) for 2 h. Finally, the samples were centrifuged for 30 min at 21,000 *g,* 4°C and the supernatant with the solubilized proteins was subjected to an affinity chromatography to purify VraS-H6x. Approximately 800 μl of a Ni Sepharose® 6 Fast Flow resin (GE Healthcare Life Sciences) was used and the procedure was carried out at 4 °C. First, the column was washed with 10 column volumes (CV) of cold distilled H_2_O followed by washing with 10 CV of Binding Buffer (Phosphate Buffer 2 or 10 mM HEPES, pH 7.5, supplemented with 0.1% DDM, 20 mM imidazole). VraS binding to the resin was carried out in batch: the resin was placed in a conical tube and solubilized fractions diluted 5 times were added to obtain a 0.1% detergent concentration. The ionic strength was increased, bringing the final concentration of NaCl to 150 mM, and the proteins were incubated with the resin for 2 h with gentle shaking. The resin was then packed in a disposable plastic column with a filter to carry out the purification by gravity and the elution from it was collected (flow-through). The resin was washed 3 times with 4 CV each, of a solution of Phosphate Buffer 2, 0.01% DDM or 10 mM HEPES, pH 7.5, 150 mM NaCl, 0.01% DDM, supplemented with increasing concentrations of imidazole (50, 75 and 100 mM). The VraS protein was eluted using 1 CV of buffer with increasing concentrations of imidazole (250, 300, 350 and 400 mM) and finished with two washes of 1.5 CV of Elution Buffer (Phosphate Buffer 2, 0.01% DDM, 500 mM imidazole or 10 mM HEPES, pH 7.5, 150 mM NaCl, 0.01% DDM, 500 mM imidazole).

### Circular Dichroism characterization of VraS

The CD spectrum of VraS was obtained by preparing a 10 μM protein solution in 10 mM sodium phosphate pH 7.4, 155 mM NaCl, 3 mM KCl, 0.01% DDM. The CD spectrum was recorded from 200 to 260 nm using a Jasco J-810 instrument (path length of the cuvette was 0.1 cm).

### Incorporation of VraS into Liposomes

The purified VraS fractions were collected (approximately 4-5 ml), supplemented with glycerol to a final concentration of 10%, and subjected to a dialysis process (10 kDa cut-off membrane; Sigma) against 500 ml of a 10 mM HEPES solution, pH 7.5, 2 mM DTT, 10% glycerol (3 changes; 4-12 h incubation each). The dialyzed fractions were centrifuged for 30 min at 18,000 *g* in a Sorvall Legend Micro 21R refrigerated centrifuge (Thermo Scientific) and the non-precipitated fraction was quantified. *E. coli* lipids (*E. coli* Polar Lipid Extract, Avanti Polar Lipids) were used to generate the liposomes. For each gram of purified protein, 20 g of lipids (1:20 protein:lipid ratio) were weighed and dissolved in 20 mM KH_2_PO_4_, pH 7 (1 ml for each gram of lipids) supplemented with n-octyl-β-D-glucopyranoside (1.5 g of detergent for each g of lipids). Once a clear solution was obtained, it was placed in a 3 kDa cut-off dialysis bag (Sigma) and dialysis was performed against 20 mM KH_2_PO_4_ solution, pH 7 (300 times greater than the volume of lipids) for 12 h at 4 °C, followed by a second dialysis against the same solution (150 times the lipid volume) for 6 h at 4 °C. The dialyzed lipid solution was subjected to three rounds of freezing in liquid nitrogen and thawing at room temperature. DDM detergent was then added at a concentration of 0.58% w/v and the purified protein (1:20 protein-lipid ratio) was added. The mixture was stirred gently for 30 min, and then incubated for 8 h at 4 °C with gentle shaking in the presence of Biobeads (Biorad; 15 mg of Biobeads for each gram of DDM). Afterward, fresh Biobeads were added and incubated for 12 additional hours at 4 °C with gentle shaking. Finally, a new portion of Biobeads was added and incubated for 4 h at room temperature with gentle shaking. Once this process was finished, the Biobeads were removed and the proteoliposome solution was aliquoted and frozen in liquid nitrogen for storage at –80 °C.

### Incubation of VraS in proteoliposomes with the vancomycin-derived photoprobe

VraS-H6x proteoliposomes (50-µL aliquotes) were placed in a 96-well microplate, and VPP was added at different concentrations (0, 6.5, 12.5, 25, 37.5, 50, 75, 100 and 125 µM). The plate was incubated for 1 h at room temperature protected from light and then placed on ice and irradiated for 2 h with a UV lamp (λ = 365 nm; Spectronics ENF-240C Handheld UV Lamp). After irradiation, the protein loading buffer solution was added to each well and the samples were resolved on a 4-12% NuPAGE gel and visualized by Western blot with a His-tag specific antibody (HRP conjugated) to detect VraS-H6x and Strep-Tactin-HRP to detect the biotin label in VPP. The intensity of the bands in the Western blot were quantified using the Gel Analyzer software; the intensity of the band obtained for the VPP-labelled VraS (Strep-Tactin-HRP blot) was relativized to the intensity of the band corresponding to the protein VraS-H6x (anti-His-tag blot), and the ratio STV/H6X was calculated for each of the conditions tested. Subsequently, the values were normalized and the affinity constant (*K*_D_) was determined with the GraphPad software, using the Hill equation for specific binding:

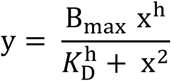

### Competition of vancomycin with VPP-binding to VraS

VraS-H6x proteoliposomes (50-µL aliquotes) were placed in a 96-well microplate wells. Different concentrations of vancomycin (0, 0.5, 1.25, 2.5, 3.75, 12.5, 25, 50 mM) were added to each well. The plate was incubated for 1 h at room temperature protected from light and then the vancomycin-derived photoprobe (VPP) was added to each well at a final concentration of 50 µM, and incubated for 1 h at room temperature, protected from light. The plate was then placed on ice and irradiated for 2 h with a UV lamp (λ = 365 nm; Spectronics ENF-240C Handheld UV Lamp). After irradiation, the protein loading buffer solution was added to each well and the samples were resolved on a 4-12% NuPAGE gel and visualized by Western blot with a His-tag specific antibody (HRP conjugated) to detect VraS-H6x and streptavidin-HRP to detect the biotin label in VPP. The intensity of the bands in the Western blot were quantified using the Gel Analyzer software; the intensity of the band obtained for the VPP-labelled VraS (streptavidin-HRP blot) was relativized to the intensity of the band corresponding to the protein VraS-H6x (anti-His-tag blot), and the ratio STV/H6X was calculated for each of the conditions tested. Subsequently, the values were normalized, and the experimental data were fitted using Graph Pad to determine the vancomycin IC50 using the following equation:

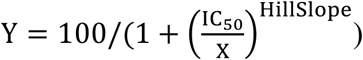

### Purification of VPP-labelled VraS

Membranes over-expressing the VraS protein were diluted to a total protein concentration of 20 mg/ml in a volume of approximately 20 ml. The sample was divided into two parts in conical tubes, one aliquot was supplemented with VPP to a final concentration of 50 μM (sample called VraS+VPP) and the other was supplemented with the same volume of DMSO (sample called VraS). Both aliquots were incubated at room temperature for 1.5 h with gentle shaking, protected from light. The contents of each conical tube were then transferred to a 6-well plate; approximately 7 ml were placed in each well. The plate was placed on ice and irradiated at 365 nm for 3 h with very gentle shaking.

After the incubation with VPP and irradiation, each one of the VraS-containing membrane preparations was incubated with 0.5% DDM in 10 mM HEPES, pH 7.5, 300 mM NaCl (approximate final volume of 40 ml). The solubilization of VraS-H6x was carried out by incubation with DDM at 4 °C for 3 h, with gentle shaking. Finally, the samples were centrifuged at 21,000 *g* for 30 min at 4 °C and the soluble fraction was subjected to an affinity chromatography in a Ni Sepharose resin.

The first step of affinity chromatography purification was performed using a Ni Sepharose® 6 Fast Flow resin (GE Healthcare). Approximately 1.5 ml of resin was used for every 40 ml of solubilized protein. In a first instance, the column was equilibrated by performing 2 washes with 5 column volumes (CV) of distilled water and then 5 washes with 10 CV of equilibration solution (10 mM HEPES, pH 7.5, 0.5% DDM, NaCl 300 mM, 20 mM imidazole). The solubilized fraction was diluted by addition of an equal volume of 10 mM HEPES buffer, pH 7.5, 300 mM NaCl, thus diluting the detergent concentration to 0.25%; imidazole was added in a final concentration of 20 mM and incubation was carried out with the equilibrated Ni Sepharose® 6 Fast Flow resin for 2 h with gentle shaking at 4 °C. The resin was then packed in a disposable affinity chromatography column with a filter (gravity flow) while the Flow-Through (FT), that is, the unbound fraction, was recirculated at least twice. Subsequently, washings were performed with: 3 CV of 10 mM HEPES, pH 7.5, 0.1% DDM, 300 mM NaCl, 50 mM imidazole; then 3 CV of 10 mM HEPES, pH 7.5, 0.05% DDM, 200 mM NaCl, 75 mM imidazole; and finally, 3 CV of 10 mM HEPES, pH 7.5, 0.025% DDM, 150 mM NaCl, 100 mM imidazole. Protein elution was performed with 1 CV of 10 mM HEPES, pH 7.5, 0.01% DDM, 150 mM NaCl, and increasing concentrations of imidazole (250, 300, 350, and 400 mM). Finally, a wash was performed with 4 CV of 10 mM HEPES, pH 7.5, 0.01% DDM, 150 mM NaCl, and 500 mM imidazole. Each eluted protein fraction (denominated at this point VraS or VraS+VPP) was supplemented with glycerol (10% final concentration) for storage at 4°C until subsequent treatment.

The VraS-VPP adduct was then purified from the VraS-H6x purified fraction (VraS+VPP) by affinity chromatography with a Streptavidin resin. Imidazole was removed from the purified VraS+VPP sample by dialysis against a 5X volume of 10 mM HEPES, pH 7.5, 10% glycerol, 150 mM NaCl. Purified fractions were placed in a 14 kDa cut-off dialysis membrane (Sigma-Aldrich) and dialyzed three times for (4-10 h), placing the dialysis bag in a new solution each time. Once the dialysis was finished, the VraS+VPP sample was centrifuged at 21,000 *g* for 20 min at 4 °C and the soluble fraction (approximately 2 ml) was incubated with 100 μL of a Streptavidin resin (streptavidin agarose conjugated, Merck) previously washed (with 10 CV of distilled water) and equilibrated. (with 20 CV of 10 mM HEPES, pH 7.5, 150 mM NaCl, 0.01% DDM and

10% glycerol). Incubation was carried out for 2 h with gentle shaking at room temperature. The resin was packed on a Micro Bio-Spin (Bio-rad) chromatography column by centrifugation (3 min at 7,000 *g*), as the Flow-Through (FT) was collected. The resin was washed 3 times with 3 CV of a 10 mM HEPES, pH 7.5, 150 mM NaCl, 0.01% DDM and 10% glycerol. The elution of the VraS-VPP adduct was performed after incubating the resin for 30 min with 100 μL of 100 mM Tris-HCl, pH 8.5, 4 mg/ml Biotin, 0.1% RapiGest and then heating the sample for 15 min at 50 °C prior to centrifuging for 3 min at 7,000 *g*. A final elution was performed with 100 μL of 100 mM Tris-HCl, pH 8.5, 4 mg/ml biotin, 2% SDS, incubating the resin with the solution for 30 min and subsequently heating for 15 min at 70 °C.

### VraS Autophosphorylation Assays

VraS proteoliposome preparations stored at-80 °C were thawed at 4 °C before use. The proteoliposomes (amount necessary to reach a final concentration of total protein of 2.5 µM) were equilibrated with the phosphorylation solution (50 mM Tris-HCl pH 7.4, 2.5 mM KCl, 1 mM MgCl_2_), with 50 µM ATP, in a final volume of 9.84 µL. The reaction was started by adding 0.16 µL of [32P]-γ-ATP (250 µCi, 10 µM), and incubated at 37 °C for 0, 15, 30 and 60 min. Subsequently, the reaction was stopped by adding 5X protein loading buffer and heating at 65 °C for 5 min. The protein was resolved in a 12% SDS-PAGE gel which was then exposed to an autoradiographic plate (Fuji imaging plate, Fujifilm) and the phosphorylated bands were visualized in a Typhoon™ FLA 7000 Scanner (General Electric). Subsequently, that same gel was stained with Coomassie Brilliant Blue to visualize the proteins. The gels were analyzed with the Gel Analyzer software, the intensities of the bands in the autoradiography were quantified and relativized to the intensity of the bands in the gel visualized by Coomassie Brilliant Blue staining.

## Supporting information

The Supporting Information File 2025_Antinori et al_Supp Info_Plos Pathogens.pdf contains the following information:

**Determination of the intact mass of VraS**. Protocol for Determination of the intact mass of VraS.

In-gel trypsin and chymotrypsin digestion of VraS and peptide extraction. Protocol for In-gel trypsin and chymotrypsin digestion of VraS and peptide extraction.

**In-solution VraS digestion.** Protocol for In-solution VraS digestion.

**Peptide identification.** Protocol for Peptide identification.

**S1 Fig. Transmembrane protein topology prediction for VraS. A.** Transmembrane α-helices in VraS were predicted using DeepTMHMM (https://dtu.biolib.com/DeepTMHMM). Red: membrane. Blue: outside/extracellular.

Pink: inside/cytoplasmic.

**S2 Fig. Non-cell wall active antibiotics do not activate the VraTSR system.** Activation of the system was monitored as a function of time in *S. aureus* RN4220/pCN52::*opvra* cultures grown in the absence or the presence of increasing concentrations of kanamycin (KN), rifampicin (RIF), linezolid (LNZ), Ciprofloxacin (CPR), Daptomycin (DAP) and polymyxin B (POLB). The graphs show the change in relative fluorescence after a two-hour incubation with the antibiotic, expressed as Δ(F/OD)_2h_; antibiotic concentrations ranged from 1x MIC to 10x MIC of each antibiotic. As a control, the effect of the solvent in which the antibiotic was dissolved was evaluated (no antibiotic). In *S. aureus* RN4220/pCN52::*opvra* cultures, the intensity of GFP fluorescence emission was recorded at λ_em_ = 528 nm, upon excitation at 485 nm and OD_600nm_ was measured simultaneously; at each time point, F/OD_600nm_ was calculated. F/OD_600nm_ was also calculated for the control strain *S. aureus* RN4220/pCN52 (autofluorescence) and was subtracted from the relative fluorescence of the reporter strain *S. aureus* RN4220/pCN52::*opvra*, to give (F/OD)_t_ = (F/ODpCN52::*opvra*)_t_ – (F/ODpCN52)_t_. The graphs show Δ(F/OD)_2h_ = (F/ODpCN52::*opvra* – F/ODpCN52)_2h_ – (F/ODpCN52::*opvra* – F/ODpCN52)_t=0h_.The results represent the means of two experiments, and the error bars correspond to the standard deviations (SD).

**S3 Fig. Activation of the VraTSR system as a function of time by vancomycin and the vancomycin-derived photoprobe (VPP).** The graphs show the change in relative fluorescence over time, expressed as Δ(F/OD)_t_, upon incubation with 5 μg/ml vancomycin or VPP. As a control, the effect of DMSO, the solvent in which the photoprobe was dissolved, was evaluated. In *S. aureus* RN4220/pCN52::*opvra* cultures, the intensity of GFP fluorescence emission was recorded at λ_em_ = 528 nm, upon excitation at 485 nm and OD_600nm_ was measured simultaneously; at each time point, F/OD_600nm_ was calculated. F/OD_600nm_ was also calculated for the control strain *S. aureus* RN4220/pCN52 (autofluorescence) and was subtracted from the relative fluorescence of the reporter strain *S. aureus* RN4220/pCN52::*opvra*, to give (F/OD)_t_ = (F/ODpCN52::*opvra*)_t_ – (F/ODpCN52)_t_. The graphs show Δ(F/OD)_t_ = (F/ODpCN52::*opvra* – F/ODpCN52)_t_ – (F/ODpCN52::*opvra* – F/ODpCN52)_t=0h_.The results represent the means of six experiments, and the error bars correspond to the standard deviations (SD).

**S4 Fig. VraS Dimer en DDM micelles and AlphaFold3 Model. A.** VraS dimer in DDM micelles. Size exclusion chromatography showed that VraS eluted at 1.87 ml in a 3 ml column (Superdex 200 5/150, Cytiva). According to the calibration curve of the column (gel filtration standard Bio-Rad #1511901), the elution volume of VraS corresponded to a molecular weight of 112 kDa, which would correspond to a dimer of VraS (2 x 40 kDa) + DDM micelle. **B.** AlphaFold3 predicted VraS dimer (one monomer colored yellow and the other light blue), showing the hydrophobic residues in the predicted extracellular loop peptide in red sticks. **C.** Hydrophobic surface of the AlphaFold3-predicted VraS dimer.

**S5 Fig. Duplicate Western blots of the ones shown in Fig 7**.

**S6 Fig. Intact mass spectrum of the VraS and VraS+VPP purified samples.** Spectra obtained on an Agilent 6546 LC/Q-TOF equipment of the proteins purified using a Ni Sepharose® 6 Fast Flow resin. A. VraS protein. B. VraS protein purified after incubation with VPP—expected molecular mass for VraS-H6x: 40,867.37 Da. VPP is 1923 Da. VraS-H6x-VPP expected mass: 42,790.37 Da.

**S7 Fig. Coverage for VraS-H6x peptides identified from in-solution chymotrypsin digestion of VraS and VraS+VPP samples purified by affinity chromatography on a Ni Sepharose® 6 Fast Flow resin.** Approximately 100 μg of VraS and 50 μg of VraS+VPP were digested with chymotrypsin in solution overnight, using a 1:100 chymotrypsin:protein ratio. Digestion was analyzed using a UHPLC Agilent 1290 Infinity II LC System coupled to a Q-TOF 6550 spectrometer (Agilent). Red bold indicates an MS/MS sequenced peptide with high confidence, identified using PEAK Studio software. Samples identified VraS-H6x with > 99.9% confidence. No VPP-bound peptides could be identified. The N-terminal (residues 1-77) chymotryptic peptides could not be identified. A. VraS sample. B. VraS+VPP sample. C. Chymotrypsin cleavage sites in the N-terminal region of VraS. Prediction made with PeptideCutter software. The predicted transmembrane helices are shown with a grey box, according to the topology predictions made with the HMMTOP, TMpred, Octupus, and TMHMM software.

**S8 Fig. Coverage for VraS-H6x peptides identified from in-solution trypsin digestion of VraS and VraS+VPP samples purified by affinity chromatography on a Ni Sepharose® 6 Fast Flow resin.** The purified VraS and VraS+VPP samples were precipitated with acetone and re-solubilized in 8 M urea. Afterwards, approximately 100 μg of VraS and VraS+VPP were digested with chymotrypsin in solution overnight, using a 1:100 chymotrypsin:protein ratio. Digestion was analyzed using a UHPLC Agilent 1290 Infinity II LC System coupled to a Q-TOF 6550 spectrometer (Agilent). Red bold indicates an MS/MS sequenced peptide with high confidence, identified using PEAK Studio software. Samples identified VraS-H6x with > 99.9% confidence. No VPP-bound peptides could be identified. The N-terminal (residues 1-70) tryptic peptides could not be identified. A. VraS sample. B. VraS+VPP sample.

**S9 Fig. Coverage for VraS-H6x peptides identified from in-solution chymotrypsin digestion of the VraS-VPP sample purified by affinity chromatography on a Streptavidin resin.** The peptides identified after chymotrypsin treatment using PEAK Studio software are shown in red. Approximately 50 μg of VraS-VPP were digested with chymotrypsin, 1:100 ON ratio. Digestion was analyzed by UHPLC 1290 Infinity II (Agilent) coupled to a Q-TOF 6550 spectrometer (Agilent).

**S10 Fig. Saturation Transfer Difference NMR spectra of a VraS-vancomycin mixture, compared to VraS and vancomycin alone. Complete spectra corresponding to Fig 9**. STD were carried out saturating at 0.75 ppm (on resonance). Buffer was 20 mM sodium phosphate pH 7.4, 280 mM NaCl, 6 mM *KCl,* 0.01% DDM, 10% deuterium oxide. From top to bottom: ^1^H NMR spectrum of vancomycin; ^1^H NMR spectrum of the vancomycin-VraS mixture used in the STD experiment (20 µM full-length VraS with 400 µM vancomycin solution); STD of 20 µM full-length VraS with 400 µM vancomycin; STD of 400 µM vancomycin; STD of 20 µM full-length VraS.

**S11 Fig. Saturation Transfer Difference NMR spectra of a VraS-ampicillin mixture, compared to VraS and ampicillin alone. Complete spectra corresponding to Fig 10**. STD were carried out saturating at 0.75 ppm (on resonance). Buffer was 20 mM sodium phosphate pH 7.4, 280 mM NaCl, 6 mM *KCl,* 0.01% DDM, 10% deuterium oxide. From top to bottom: ^1^H NMR spectrum of ampicillin; ^1^H NMR spectrum of the ampicillin-VraS mixture used in the STD experiment (20 µM full-length VraS with 400 µM ampicillin solution); STD of 20 µM full-length VraS with 400 µM ampicillin; STD of 400 µM ampicillin; STD of 20 µM full-length VraS.

**S12 Fig. Saturation Transfer Difference NMR assays of a kanamycin-VraS mixture did not show evidence of an interaction.** ^1^H NMR spectrum of 400 µM kanamycin (top), and Saturation Transfer Difference NMR spectra, saturating at 0.75 ppm, of 20 µM full-length VraS with 400 µM kanamycin (middle) and of 400 µM kanamycin (bottom). The peaks inside the colored rectangles, in the ^1^H NMR spectrum of the kanamycin, correspond to the protons of kanamycin highlighted in the structure with a circle of the same color, and were not detected in the STD experiment of the kanamycin-VraS mixture. Buffer was 20 mM sodium phosphate pH 7.4, 280 mM NaCl, 6 mM KCl, 0.01% DDM, 10% deuterium oxide.

## Acknowledgments

M.B.A. was a Ph.D. Fellow of CONICET. M.B.A. was recipient of a PROLAB Fellowship from ASBMB. I.P.S. and L.I.L. are Staff members of CONICET. We thank Andrea Coscia, Alejandro Gago and Dr. Rodolfo M. Rasia from the Argentinean Structural Biology and Metabolomics Platform (PLABEM, IBR-UNR-CONICET) for their support on NMR experiments and Marina Perozzi from IBR-UNR-CONICET for her assistance on research activities.

## Author contributions

Conceptualization and Project Administration, L.I.L; Investigation, M.B.A., D.S., K.K., I.P.S. and M.D.P.; Methodology, L.I.L., I.P.S., D.S.; Visualization, M.B.A., I.P.S. and L.I.L.; Writing – Original Draft Preparation, L.I.L., I.P.S. and D.S.; Review and Editing, L.I.L., D.S, K.K., G.D.W., M.B.A., I.P.S. and M.D.P.; Funding Acquisition and Supervision, L.I.L. and G.D.W.

## References

1. Grundmann H, Aires-de-Sousa M, Boyce J, Tiemersma E. Emergence and resurgence of meticillin-resistant Staphylococcus aureus as a public-health threat. Lancet. 2006;368(9538):874-85. Epub 2006/09/05. doi: 10.1016/S0140-6736(06)68853-3. PubMed PMID: 16950365.

2. Tong SY, McDonald MI, Holt DC, Currie BJ. Global implications of the emergence of community-associated methicillin-resistant Staphylococcus aureus in Indigenous populations. Clin Infect Dis. 2008;46(12):1871–8. Epub 2008/05/09. doi: 10.1086/588301. PubMed PMID: 18462175.

3. Poulakou G, Lagou S, Tsiodras S. What’s new in the epidemiology of skin and soft tissue infections in 2018? Current opinion in infectious diseases. 2019;32(2):77–86. doi: 10.1097/QCO.0000000000000527. PubMed PMID: 30664027.

4. Woodford N, Livermore DM. Infections caused by Gram-positive bacteria: a review of the global challenge. J Infect. 2009;59 Suppl 1:S4–16. Epub 2009/09/22. doi: 10.1016/S0163-4453(09)60003-7. PubMed PMID: 19766888.

5. Boucher HW, Talbot GH, Bradley JS, Edwards JE, Gilbert D, Rice LB, et al. Bad bugs, no drugs: no ESKAPE! An update from the Infectious Diseases Society of America. Clin Infect Dis. 2009;48(1):1–12. Epub 2008/11/28. doi: 10.1086/595011. PubMed PMID: 19035777.

6. Gajdacs M. The Continuing Threat of Methicillin-Resistant Staphylococcus aureus. Antibiotics. 2019;8(2). doi: 10.3390/antibiotics8020052. PubMed PMID: 31052511; PubMed Central PMCID: PMC6627156.

7. Guzman-Blanco M, Mejia C, Isturiz R, Alvarez C, Bavestrello L, Gotuzzo E, et al. Epidemiology of methicillin-resistant Staphylococcus aureus (MRSA) in Latin America. Int J Antimicrob Agents. 2009;34(4):304–8. Epub 2009/07/25. doi: 10.1016/j.ijantimicag.2009.06.005. PubMed PMID: 19625169.

8. Rodriguez-Noriega E, Seas C, Guzman-Blanco M, Mejia C, Alvarez C, Bavestrello L, et al. Evolution of methicillin-resistant Staphylococcus aureus clones in Latin America. Int J Infect Dis. 2010;14(7):e560–6. Epub 2010/01/06. doi: 10.1016/j.ijid.2009.08.018. PubMed PMID: 20047848.

9. Livermore DM. Has the era of untreatable infections arrived? J Antimicrob Chemother. 2009;64 Suppl 1:i29–36. Epub 2009/08/19. doi: 10.1093/jac/dkp255. PubMed PMID: 19675016.

10. The antibiotic alarm. Nature. 2013;495(7440):141. Epub 2013/03/16. doi: 10.1038/495141a. PubMed PMID: 23495392.

11. Organization WH. Addressing the crisis in antibiotic development 2020. Available from: https://www.who.int/news/item/09-07-2020-addressing-the-crisis-in-antibiotic-development.

12. Kuroda M, Kuwahara-Arai K, Hiramatsu K. Identification of the up-and down-regulated genes in vancomycin-resistant Staphylococcus aureus strains Mu3 and Mu50 by cDNA differential hybridization method. Biochem Biophys Res Commun. 2000;269(2):485–90. Epub 2000/03/10. doi: 10.1006/bbrc.2000.2277. PubMed PMID: 10708580.

13. Kuroda M, Kuroda H, Oshima T, Takeuchi F, Mori H, Hiramatsu K. Two-component system VraSR positively modulates the regulation of cell-wall biosynthesis pathway in Staphylococcus aureus. Mol Microbiol. 2003;49(3):807–21. Epub 2003/07/17. doi: 10.1046/j.1365-2958.2003.03599.x. PubMed PMID: 12864861.

14. Gardete S, Wu SW, Gill S, Tomasz A. Role of VraSR in antibiotic resistance and antibiotic-induced stress response in Staphylococcus aureus. Antimicrob Agents Chemother. 2006;50(10):3424–34. Epub 2006/09/29. doi: 10.1128/AAC.00356-06. PubMed PMID: 17005825; PubMed Central PMCID: PMC1610096.

15. Utaida S, Dunman PM, Macapagal D, Murphy E, Projan SJ, Singh VK, et al. Genome-wide transcriptional profiling of the response of Staphylococcus aureus to cell-wall-active antibiotics reveals a cell-wall-stress stimulon. Microbiology (Reading). 2003;149(Pt 10):2719–32. Epub 2003/10/03. doi: 10.1099/mic.0.26426-0. PubMed PMID: 14523105.

16. Dengler V, Meier PS, Heusser R, Berger-Bachi B, McCallum N. Induction kinetics of the Staphylococcus aureus cell wall stress stimulon in response to different cell wall active antibiotics. BMC Microbiol. 2011;11:16. Epub 2011/01/22. doi: 10.1186/1471-2180-11-16. PubMed PMID: 21251258; PubMed Central PMCID: PMCPMC3032642.

17. Huber J, Donald RG, Lee SH, Jarantow LW, Salvatore MJ, Meng X, et al. Chemical genetic identification of peptidoglycan inhibitors potentiating carbapenem activity against methicillin-resistant Staphylococcus aureus. Chem Biol. 2009;16(8):837–48. Epub 2009/09/01. doi: 10.1016/j.chembiol.2009.05.012. PubMed PMID: 19716474.

18. Katayama Y, Murakami-Kuroda H, Cui L, Hiramatsu K. Selection of heterogeneous vancomycin-intermediate Staphylococcus aureus by imipenem. Antimicrob Agents Chemother. 2009;53(8):3190–6. Epub 2009/05/20. doi: 10.1128/AAC.00834-08. PubMed PMID: 19451283; PubMed Central PMCID: PMC2715608.

19. Kato Y, Suzuki T, Ida T, Maebashi K. Genetic changes associated with glycopeptide resistance in Staphylococcus aureus: predominance of amino acid substitutions in YvqF/VraSR. J Antimicrob Chemother. 2010;65(1):37–45. Epub 2009/11/06. doi: 10.1093/jac/dkp394. PubMed PMID: 19889788; PubMed Central PMCID: PMC2800785.

20. Yoo JI, Kim JW, Kang GS, Kim HS, Yoo JS, Lee YS. Prevalence of amino acid changes in the yvqF, vraSR, graSR, and tcaRAB genes from vancomycin intermediate resistant Staphylococcus aureus. J Microbiol. 2013;51(2):160–5. Epub 2013/04/30. doi: 10.1007/s12275-013-3088-7. PubMed PMID: 23625215.

21. Sieradzki K, Tomasz A. Alterations of cell wall structure and metabolism accompany reduced susceptibility to vancomycin in an isogenic series of clinical isolates of Staphylococcus aureus. J Bacteriol. 2003;185(24):7103–10. Epub 2003/12/04. doi: 10.1128/JB.185.24.7103-7110.2003. PubMed PMID: 14645269; PubMed Central PMCID: PMCPMC296238.

22. Sieradzki K, Tomasz A. Gradual alterations in cell wall structure and metabolism in vancomycin-resistant mutants of Staphylococcus aureus. J Bacteriol. 1999;181(24):7566–70. Epub 1999/12/22. doi: 10.1128/JB.181.24.7566-7570.1999. PubMed PMID: 10601215; PubMed Central PMCID: PMCPMC94215.

23. Boyle-Vavra S, Yin S, Daum RS. The VraS/VraR two-component regulatory system required for oxacillin resistance in community-acquired methicillin-resistant Staphylococcus aureus. FEMS Microbiol Lett. 2006;262(2):163–71. Epub 2006/08/23. doi: 10.1111/j.1574-6968.2006.00384.x. PubMed PMID: 16923071.

24. Mehta S, Cuirolo AX, Plata KB, Riosa S, Silverman JA, Rubio A, et al. VraSR two-component regulatory system contributes to mprF-mediated decreased susceptibility to daptomycin in in vivo-selected clinical strains of methicillin-resistant Staphylococcus aureus. Antimicrob Agents Chemother. 2012;56(1):92–102. Epub 2011/10/12. doi: 10.1128/AAC.00432-10. PubMed PMID: 21986832; PubMed Central PMCID: PMCPMC3256076.

25. Qureshi NK, Yin S, Boyle-Vavra S. The role of the Staphylococcal VraTSR regulatory system on vancomycin resistance and vanA operon expression in vancomycin-resistant Staphylococcus aureus. PLoS One. 2014;9(1):e85873. Epub 2014/01/24. doi: 10.1371/journal.pone.0085873. PubMed PMID: 24454941; PubMed Central PMCID: PMCPMC3893269.

26. Jordan S, Junker A, Helmann JD, Mascher T. Regulation of LiaRS-dependent gene expression in bacillus subtilis: identification of inhibitor proteins, regulator binding sites, and target genes of a conserved cell envelope stress-sensing two-component system. J Bacteriol. 2006;188(14):5153–66. Epub 2006/07/04. doi: 10.1128/JB.00310-06. PubMed PMID: 16816187; PubMed Central PMCID: PMCPMC1539951.

27. Eldholm V, Gutt B, Johnsborg O, Bruckner R, Maurer P, Hakenbeck R, et al. The pneumococcal cell envelope stress-sensing system LiaFSR is activated by murein hydrolases and lipid II-interacting antibiotics. J Bacteriol. 2010;192(7):1761–73. Epub 2010/02/02. doi: 10.1128/JB.01489-09. PubMed PMID: 20118250; PubMed Central PMCID: PMCPMC2838051.

28. Martinez B, Zomer AL, Rodriguez A, Kok J, Kuipers OP. Cell envelope stress induced by the bacteriocin Lcn972 is sensed by the Lactococcal two-component system CesSR. Mol Microbiol. 2007;64(2):473–86. Epub 2007/05/12. doi: 10.1111/j.1365-2958.2007.05668.x. PubMed PMID: 17493129.

29. Jordan S, Hutchings MI, Mascher T. Cell envelope stress response in Gram-positive bacteria. FEMS Microbiol Rev. 2008;32(1):107–46. Epub 2008/01/05. doi: 10.1111/j.1574-6976.2007.00091.x. PubMed PMID: 18173394.

30. Boyle-Vavra S, Yin S, Jo DS, Montgomery CP, Daum RS. VraT/YvqF is required for methicillin resistance and activation of the VraSR regulon in Staphylococcus aureus. Antimicrob Agents Chemother. 2013;57(1):83–95. Epub 2012/10/17. doi: 10.1128/AAC.01651-12. PubMed PMID: 23070169; PubMed Central PMCID: PMC3535960.

31. McCallum N, Meier PS, Heusser R, Berger-Bachi B. Mutational analyses of open reading frames within the vraSR operon and their roles in the cell wall stress response of Staphylococcus aureus. Antimicrob Agents Chemother. 2011;55(4):1391–402. Epub 2011/01/12. doi: 10.1128/AAC.01213-10. PubMed PMID: 21220524; PubMed Central PMCID: PMCPMC3067146.

32. Belcheva A, Golemi-Kotra D. A close-up view of the VraSR two-component system. A mediator of Staphylococcus aureus response to cell wall damage. J Biol Chem. 2008;283(18):12354–64. Epub 2008/03/11. doi: 10.1074/jbc.M710010200. PubMed PMID: 18326495.

33. Fernandes PB, Reed P, Monteiro JM, Pinho MG. Revisiting the Role of VraTSR in Staphylococcus aureus Response to Cell Wall-Targeting Antibiotics. J Bacteriol. 2022;204(8):e0016222. Epub 2022/07/22. doi: 10.1128/jb.00162-22. PubMed PMID: 35862765; PubMed Central PMCID: PMCPMC9380581.

34. Juan C, Torrens G, Gonzalez-Nicolau M, Oliver A. Diversity and regulation of intrinsic beta-lactamases from non-fermenting and other Gram-negative opportunistic pathogens. FEMS Microbiol Rev. 2017;41(6):781–815. Epub 2017/10/14. doi: 10.1093/femsre/fux043. PubMed PMID: 29029112.

35. Wang Q, Marchetti R, Prisic S, Ishii K, Arai Y, Ohta I, et al. A Comprehensive Study of the Interaction between Peptidoglycan Fragments and the Extracellular Domain of Mycobacterium tuberculosis Ser/Thr Kinase PknB. Chembiochem. 2017;18(21):2094–8. Epub 2017/08/30. doi: 10.1002/cbic.201700385. PubMed PMID: 28851116; PubMed Central PMCID: PMCPMC6261334.

36. Dik DA, Dominguez-Gil T, Lee M, Hesek D, Byun B, Fishovitz J, et al. Muropeptide Binding and the X-ray Structure of the Effector Domain of the Transcriptional Regulator AmpR of Pseudomonas aeruginosa. J Am Chem Soc. 2017;139(4):1448–51. Epub 2017/01/13. doi: 10.1021/jacs.6b12819. PubMed PMID: 28079369; PubMed Central PMCID: PMCPMC5436579.

37. Tan S, Cho K, Nodwell JR. A defect in cell wall recycling confers antibiotic resistance and sensitivity in Staphylococcus aureus. J Biol Chem. 2022;298(10):102473. Epub 2022/09/12. doi: 10.1016/j.jbc.2022.102473. PubMed PMID: 36089064; PubMed Central PMCID: PMCPMC9547203.

38. Arthur M, Molinas C, Courvalin P. The VanS-VanR two-component regulatory system controls synthesis of depsipeptide peptidoglycan precursors in Enterococcus faecium BM4147. J Bacteriol. 1992;174(8):2582–91. doi: 10.1128/jb.174.8.2582-2591.1992. PubMed PMID: 1556077; PubMed Central PMCID: PMCPMC205897.

39. Hutchings MI, Hong HJ, Buttner MJ. The vancomycin resistance VanRS two-component signal transduction system of Streptomyces coelicolor. Mol Microbiol. 2006;59(3):923–35. Epub 2006/01/20. doi: 10.1111/j.1365-2958.2005.04953.x. PubMed PMID: 16420361.

40. Koteva K, Hong HJ, Wang XD, Nazi I, Hughes D, Naldrett MJ, et al. A vancomycin photoprobe identifies the histidine kinase VanSsc as a vancomycin receptor. Nat Chem Biol. 2010;6(5):327–9. Epub 2010/04/13. doi: 10.1038/nchembio.350. PubMed PMID: 20383152.

41. Lockey C, Edwards RJ, Roper DI, Dixon AM. The Extracellular Domain of Two-component System Sensor Kinase VanS from Streptomyces coelicolor Binds Vancomycin at a Newly Identified Binding Site. Scientific reports. 2020;10(1):5727. Epub 2020/04/03. doi: 10.1038/s41598-020-62557-z. PubMed PMID: 32235931; PubMed Central PMCID: PMCPMC7109055.

42. Steidl R, Pearson S, Stephenson RE, Ledala N, Sitthisak S, Wilkinson BJ, Jayaswal RK. Staphylococcus aureus cell wall stress stimulon gene-lacZ fusion strains: potential for use in screening for cell wall-active antimicrobials. Antimicrob Agents Chemother. 2008;52(8):2923–5. Epub 2008/06/11. doi: 10.1128/AAC.00273-08. PubMed PMID: 18541730; PubMed Central PMCID: PMCPMC2493131.

43. Belcheva A, Verma V, Golemi-Kotra D. DNA-binding activity of the vancomycin resistance associated regulator protein VraR and the role of phosphorylation in transcriptional regulation of the vraSR operon. Biochemistry. 2009;48(24):5592–601. Epub 2009/05/08. doi: 10.1021/bi900478b. PubMed PMID: 19419158.

44. Reynolds PE. Structure, biochemistry and mechanism of action of glycopeptide antibiotics. European journal of clinical microbiology & infectious diseases: official publication of the European Society of Clinical Microbiology. 1989;8(11):943–50. Epub 1989/11/01. doi: 10.1007/BF01967563. PubMed PMID: 2532132.

45. Culp EJ, Waglechner N, Wang W, Fiebig-Comyn AA, Hsu YP, Koteva K, et al. Evolution-guided discovery of antibiotics that inhibit peptidoglycan remodelling. Nature. 2020;578(7796):582-7. Epub 2020/02/14. doi: 10.1038/s41586-020-1990-9. PubMed PMID: 32051588.

46. Xu M, Wang W, Waglechner N, Culp EJ, Guitor AK, Wright GD. Phylogeny-Informed Synthetic Biology Reveals Unprecedented Structural Novelty in Type V Glycopeptide Antibiotics. ACS Cent Sci. 2022;8(5):615–26. Epub 2022/06/02. doi: 10.1021/acscentsci.1c01389. PubMed PMID: 35647273; PubMed Central PMCID: PMCPMC9136965.

47. Koteva K, Xu M, Wang W, Fiebig-Comyn AA, Cook MA, Coombes BK, Wright GD. Synthetic Biology Facilitates Semisynthetic Development of Type V Glycopeptide Antibiotics Targeting Vancomycin-Resistant Enterococcus. J Med Chem. 2023;66(13):9006–22. Epub 2023/06/14. doi: 10.1021/acs.jmedchem.3c00633. PubMed PMID: 37315221; PubMed Central PMCID: PMCPMC10350919.

48. Lamb SS, Patel T, Koteva KP, Wright GD. Biosynthesis of sulfated glycopeptide antibiotics by using the sulfotransferase StaL. Chem Biol. 2006;13(2):171–81. Epub 2006/02/24. doi: 10.1016/j.chembiol.2005.12.003. PubMed PMID: 16492565.

49. Yim G, Wang W, Pawlowski AC, Wright GD. Trichlorination of a Teicoplanin-Type Glycopeptide Antibiotic by the Halogenase StaI Evades Resistance. Antimicrob Agents Chemother. 2018;62(12). Epub 2018/10/03. doi: 10.1128/AAC.01540-18. PubMed PMID: 30275088; PubMed Central PMCID: PMCPMC6256792.

50. Baizman ER, Branstrom AA, Longley CB, Allanson N, Sofia MJ, Gange D, Goldman RC. Antibacterial activity of synthetic analogues based on the disaccharide structure of moenomycin, an inhibitor of bacterial transglycosylase. Microbiology (Reading). 2000;146 Pt 12:3129–40. Epub 2000/12/02. doi: 10.1099/00221287-146-12-3129. PubMed PMID: 11101671.

51. Goldman RC, Gange D. Inhibition of transglycosylation involved in bacterial peptidoglycan synthesis. Curr Med Chem. 2000;7(8):801–20. Epub 2000/06/01. doi: 10.2174/0929867003374651. PubMed PMID: 10828288.

52. Halliday J, McKeveney D, Muldoon C, Rajaratnam P, Meutermans W. Targeting the forgotten transglycosylases. Biochem Pharmacol. 2006;71(7):957–67. Epub 2005/11/22. doi: 10.1016/j.bcp.2005.10.030. PubMed PMID: 16298347.

53. Sabat AJ, Tinelli M, Grundmann H, Akkerboom V, Monaco M, Del Grosso M, et al. Daptomycin Resistant Staphylococcus aureus Clinical Strain With Novel Non-synonymous Mutations in the mprF and vraS Genes: A New Insight Into Daptomycin Resistance. Front Microbiol. 2018;9:2705. Epub 2018/11/22. doi: 10.3389/fmicb.2018.02705. PubMed PMID: 30459746; PubMed Central PMCID: PMCPMC6232378.

54. Hallgren J, Tsirigos KD, Pedersen MD, Almagro Armenteros JJ, Marcatili P, Nielsen H, et al. DeepTMHMM predicts alpha and beta transmembrane proteins using deep neural networks. bioRxiv. 2022:2022.04.08.487609. doi: 10.1101/2022.04.08.487609.

55. Gutierrez S, Tyczynski WG, Boomsma W, Teufel F, Winther O. MembraneFold: Visualising transmembrane protein structure and topology. bioRxiv. 2022:2022.12.06.518085. doi: 10.1101/2022.12.06.518085.

56. Abramson J, Adler J, Dunger J, Evans R, Green T, Pritzel A, et al. Accurate structure prediction of biomolecular interactions with AlphaFold 3. Nature. 2024;630(8016):493-500. Epub 2024/05/09. doi: 10.1038/s41586-024-07487-w. PubMed PMID: 38718835; PubMed Central PMCID: PMCPMC11168924 including 63/611,674, 63/611,638 and 63/546,444 relating to predicting 3D structures of molecule complexes using embedding neural networks and generative models. All of the authors other than A.B., Y.A.K. and E.D.Z. have commercial interests in the work described.

57. Kim DJ, Forst S. Genomic analysis of the histidine kinase family in bacteria and archaea. Microbiology (Reading). 2001;147(Pt 5):1197–212. Epub 2001/04/26. doi: 10.1099/00221287-147-5-1197. PubMed PMID: 11320123.

58. Dorman G, Prestwich GD. Benzophenone photophores in biochemistry. Biochemistry. 1994;33(19):5661–73. Epub 1994/05/17. doi: 10.1021/bi00185a001. PubMed PMID: 8180191.

59. Bem AE, Velikova N, Pellicer MT, Baarlen P, Marina A, Wells JM. Bacterial histidine kinases as novel antibacterial drug targets. ACS Chem Biol. 2015;10(1):213–24. Epub 2014/12/02. doi: 10.1021/cb5007135. PubMed PMID: 25436989.

60. Hirakawa H, Kurushima J, Hashimoto Y, Tomita H. Progress Overview of Bacterial Two-Component Regulatory Systems as Potential Targets for Antimicrobial Chemotherapy. Antibiotics. 2020;9(10). Epub 2020/09/27. doi: 10.3390/antibiotics9100635. PubMed PMID: 32977461; PubMed Central PMCID: PMCPMC7598275.

61. Zschiedrich CP, Keidel V, Szurmant H. Molecular Mechanisms of Two-Component Signal Transduction. J Mol Biol. 2016;428(19):3752–75. Epub 2016/08/16. doi: 10.1016/j.jmb.2016.08.003. PubMed PMID: 27519796; PubMed Central PMCID: PMCPMC5023499.

62. Sevvana M, Vijayan V, Zweckstetter M, Reinelt S, Madden DR, Herbst-Irmer R, et al. A ligand-induced switch in the periplasmic domain of sensor histidine kinase CitA. J Mol Biol. 2008;377(2):512–23. Epub 2008/02/09. doi: 10.1016/j.jmb.2008.01.024. PubMed PMID: 18258261.

63. Wu R, Gu M, Wilton R, Babnigg G, Kim Y, Pokkuluri PR, et al. Insight into the sporulation phosphorelay: crystal structure of the sensor domain of Bacillus subtilis histidine kinase, KinD. Protein Sci. 2013;22(5):564–76. Epub 2013/02/26. doi: 10.1002/pro.2237. PubMed PMID: 23436677; PubMed Central PMCID: PMCPMC3649258.

64. Bader MW, Sanowar S, Daley ME, Schneider AR, Cho U, Xu W, et al. Recognition of antimicrobial peptides by a bacterial sensor kinase. Cell. 2005;122(3):461–72. Epub 2005/08/13. doi: 10.1016/j.cell.2005.05.030. PubMed PMID: 16096064.

65. Groisman EA, Duprey A, Choi J. How the PhoP/PhoQ System Controls Virulence and Mg(2+) Homeostasis: Lessons in Signal Transduction, Pathogenesis, Physiology, and Evolution. Microbiol Mol Biol Rev. 2021;85(3):e0017620. Epub 2021/07/01. doi: 10.1128/MMBR.00176-20. PubMed PMID: 34191587; PubMed Central PMCID: PMCPMC8483708.

66. Chamnongpol S, Cromie M, Groisman EA. Mg2+ sensing by the Mg2+ sensor PhoQ of Salmonella enterica. J Mol Biol. 2003;325(4):795–807. Epub 2003/01/01. doi: 10.1016/s0022-2836(02)01268-8. PubMed PMID: 12507481.

67. Cheung J, Hendrickson WA. Structural analysis of ligand stimulation of the histidine kinase NarX. Structure. 2009;17(2):190–201. Epub 2009/02/17. doi: 10.1016/j.str.2008.12.013. PubMed PMID: 19217390; PubMed Central PMCID: PMCPMC3749045.

68. Unden G, Nilkens S, Singenstreu M. Bacterial sensor kinases using Fe-S cluster binding PAS or GAF domains for O2 sensing. Dalton Trans. 2013;42(9):3082–7. Epub 2012/11/10. doi: 10.1039/c2dt32089d. PubMed PMID: 23138661.

69. Mascher T. Intramembrane-sensing histidine kinases: a new family of cell envelope stress sensors in Firmicutes bacteria. FEMS Microbiol Lett. 2006;264(2):133–44. Epub 2006/10/27. doi: 10.1111/j.1574-6968.2006.00444.x. PubMed PMID: 17064367.

70. Fernandez P, Porrini L, Albanesi D, Abriata LA, Dal Peraro M, de Mendoza D, Mansilla MC. Transmembrane Prolines Mediate Signal Sensing and Decoding in Bacillus subtilis DesK Histidine Kinase. mBio. 2019;10(6). Epub 2019/11/28. doi: 10.1128/mBio.02564-19. PubMed PMID: 31772055; PubMed Central PMCID: PMCPMC6879721.

71. Trajtenberg F, Graña M, Ruétalo N, Botti H, Buschiazzo A. Structural and enzymatic insights into the ATP binding and autophosphorylation mechanism of a sensor histidine kinase. J Biol Chem. 2010;285(32):24892–903. Epub 2010/05/29. doi: 10.1074/jbc.M110.147843. PubMed PMID: 20507988; PubMed Central PMCID: PMCPMC2915725.

72. Jani S, Sterzenbach K, Adatrao V, Tajbakhsh G, Mascher T, Golemi-Kotra D. Low phosphatase activity of LiaS and strong LiaR-DNA affinity explain the unusual LiaS to LiaR in vivo stoichiometry. BMC Microbiol. 2020;20(1):104. Epub 2020/05/01. doi: 10.1186/s12866-020-01796-6. PubMed PMID: 32349670; PubMed Central PMCID: PMCPMC7191749.

73. Mascher T, Zimmer SL, Smith TA, Helmann JD. Antibiotic-inducible promoter regulated by the cell envelope stress-sensing two-component system LiaRS of Bacillus subtilis. Antimicrob Agents Chemother. 2004;48(8):2888–96. Epub 2004/07/27. doi: 10.1128/AAC.48.8.2888-2896.2004. PubMed PMID: 15273097; PubMed Central PMCID: PMCPMC478541.

74. Mascher T, Helmann JD, Unden G. Stimulus perception in bacterial signal-transducing histidine kinases. Microbiol Mol Biol Rev. 2006;70(4):910–38. Epub 2006/12/13. doi: 10.1128/MMBR.00020-06. PubMed PMID: 17158704; PubMed Central PMCID: PMCPMC1698512.

75. Pootoolal J, Thomas MG, Marshall CG, Neu JM, Hubbard BK, Walsh CT, Wright GD. Assembling the glycopeptide antibiotic scaffold: The biosynthesis of A47934 from Streptomyces toyocaensis NRRL15009. Proc Natl Acad Sci U S A. 2002;99(13):8962–7. Epub 2002/06/13. doi: 10.1073/pnas.102285099. PubMed PMID: 12060705; PubMed Central PMCID: PMCPMC124406.

76. Gomez-Arrebola C, Hernandez SB, Culp EJ, Wright GD, Solano C, Cava F, Lasa I. Staphylococcus aureus susceptibility to complestatin and corbomycin depends on the VraSR two-component system. Microbiol Spectr. 2023;11(5):e0037023. Epub 2023/08/30. doi: 10.1128/spectrum.00370-23. PubMed PMID: 37646518; PubMed Central PMCID: PMCPMC10581084.

77. McCallum N, Spehar G, Bischoff M, Berger-Bachi B. Strain dependence of the cell wall-damage induced stimulon in Staphylococcus aureus. Biochim Biophys Acta. 2006;1760(10):1475–81. Epub 2006/08/08. doi: 10.1016/j.bbagen.2006.06.008. PubMed PMID: 16891058.

78. Gao M, Skolnick J. The distribution of ligand-binding pockets around protein-protein interfaces suggests a general mechanism for pocket formation. Proc Natl Acad Sci U S A. 2012;109(10):3784–9. Epub 2012/02/23. doi: 10.1073/pnas.1117768109. PubMed PMID: 22355140; PubMed Central PMCID: PMCPMC3309739.

79. Skolnick J, Zhou H. Implications of the Essential Role of Small Molecule Ligand Binding Pockets in Protein-Protein Interactions. J Phys Chem B. 2022;126(36):6853–67. Epub 2022/09/01. doi: 10.1021/acs.jpcb.2c04525. PubMed PMID: 36044742; PubMed Central PMCID: PMCPMC9484464.

80. Nair D, Memmi G, Hernandez D, Bard J, Beaume M, Gill S, et al. Whole-genome sequencing of Staphylococcus aureus strain RN4220, a key laboratory strain used in virulence research, identifies mutations that affect not only virulence factors but also the fitness of the strain. J Bacteriol. 2011;193(9):2332–5. Epub 2011/03/08. doi: 10.1128/JB.00027-11. PubMed PMID: 21378186; PubMed Central PMCID: PMCPMC3133102.

81. Charpentier E, Anton AI, Barry P, Alfonso B, Fang Y, Novick RP. Novel cassette-based shuttle vector system for gram-positive bacteria. Appl Environ Microbiol. 2004;70(10):6076–85. Epub 2004/10/07. doi: 10.1128/AEM.70.10.6076-6085.2004. PubMed PMID: 15466553; PubMed Central PMCID: PMC522135.

